# LiDAR and hyperspectral-based structured population models show future forest fire frequency may compromise forest resilience

**DOI:** 10.1101/2025.07.23.666246

**Authors:** Jessica McLean, Tommaso Jucker, Alice Rosen, Sean M. McMahon, Roberto Salguero-Gómez

**Author notes:** **Author email addresses (: contact authors)** Jessica McLean Alice Rosen Sean M. McMahon Tommaso Jucker Roberto Salguero-Gómez.

## Abstract

Forest disturbances are accelerating biodiversity loss and altering tree productivity worldwide. Post-disturbance recovery time is critical for identifying vulnerable areas and targeting conservation but varies with environmental conditions. Monitoring recovery at scale requires tracking tree dynamics, yet traditional ground-based approaches are resource-intensive. We present a pipeline to parameterise integral projection models (IPMs) using LiDAR and hyperspectral data to assess post-fire recovery across large, forested areas. Focusing on the fire-adapted *Picea mariana*, we model passage times to reproductive heights and life expectancy under different fire regimes as indicators of recovery. To do this, we combined hyperspectral-derived species maps and LiDAR-based crown heights to track individual tree survival and growth at the Caribou-Poker Creek Research Watershed (BONA) from 2017–2023. We incorporated fire history, aspect, slope, elevation, and surrounding canopy height into our models and found partial support for their expected effects on survival and growth. Once accounting for topography and competition, we estimated passage times to reproductive maturity (11-22 years). Life expectancy in the absence of fire is shortest on North-facing slopes with recent fire (579 years). Sensitivity analyses highlight fire history and aspect as key modulators of population resilience, with elevation exerting strong influence on life expectancy across all conditions. Our results demonstrate that remotely sensed IPMs can effectively quantify forest recovery at scale, revealing that in some contexts, stands of *P. mariana* may not recover between fire disturbances. We discuss the implications of these findings for resilience-based forest management and highlight both the challenges and opportunities of using LiDAR and hyperspectral data to build demographic models for forecasting forest dynamics.

## Introduction

Forest disturbances like pests, drought, and fires affect the structure, function, and composition of forests (Canelles et al., 2021; Hammond et al., 2022; Roces-Díaz et al., 2022). Predicting the future impacts of disturbance, however, is challenging due to the varied spatial extent, species-specific vulnerabilities and resilience, and the return time of the events (Sturtevant & Fortin, 2021). Further, multiple features of individual trees, species, and the environment can have a strong influence on the vulnerability of forests, requiring more complex assessment of threats as well as requiring data on individuals, even if the goal of an analysis is to infer species, community, or even biome-scale responses to disturbance (Viljur et al., 2022). Demographic analyses offer a keyway to translate the survival and growth of individual trees into the population trends that can project future forest states (Ohse et al., 2023). Demographic data on individuals is difficult to acquire over large spatial domains, further challenging our ability to link disturbances that can span regions to projections emerging from the collective response of individual trees (Rosen et al., 2025).

Remote sensing technology, including hyperspectral imagery and Light Detection and Ranging (LiDAR) have accelerated the monitoring of forest disturbances (Fassnacht et al., 2024; Cavendar-Bares, Gamon & Townsend, 2020). Indeed, repeat airborne LiDAR acquisitions can now be used to track changes in tree canopies over thousands of hectares at unprecedented spatial resolution (1-4 m^2^) (Jucker, 2022; Battison et al., 2024). While aerial laser scanning was first utilised to understand canopy-level effects of disturbances, (Cushman et al., 2021; Choi et al., 2023), improvements in individual tree detection methods (Dalponte & Coomes, 2016; Cao et al., 2023) have opened the opportunity to track trees through time (Duncanson & Dubayah, 2018; Beese et al., 2022; Battison et al., 2024). The repeatability and spatial scale of LiDAR acquisitions overcomes the major time and labour constraints of classical approaches to collect field data (Salguero-Gómez & Gamelon, 2021). Furthermore, the differences in the reflectance spectra of different tree species can be used to identify specific populations across a landscape, especially in low-diversity systems (Dalponte & Coomes, 2016; Scholl et al., 2020; Weinstein et al., 2024). The large sample sizes resulting from remote sensing approaches (*e.g.* combining laser scanning for tree measurements and hyperspectral for species mapping) offers data for analyses at the spatial scales common to many disturbance events.

Structured population models parametrised with demographic hyperspectral and LiDAR data to estimate the effect of disturbances on vital rates like survival, growth, and reproduction. Indeed, ecologists use these models to examine the effects of habitat fragmentation (Zambrano & Salguero-Gómez, 2014), pathogens (Needham et al., 2016) or climate change (Kunstler et al., 2021) on forest dynamics. In their standard version, integral projection models (IPMs hereafter; Easterling, Ellner & Dixon, 2000; Ellner & Rees, 2006), a form of structured population model, describe the temporal changes in population distribution of individuals as a function of a continuous trait that predict vital rates (Merow et al., 2014). Historically, structured population models (including IPMs) for trees have used diameter at breast height (DBH) to predict vital rates (Metcalf et al., 2009; Needham et al., 2016, 2018; Ohse et al., 2023; Salguero-Gómez et al., 2015). However, characteristics of trees measured with LiDAR, like tree height, can also predict productivity, competition, and mortality (Jucker et al., 2017; Maynard et al., 2022; Zuleta et al., 2022). Additionally, as IPMs can easily incorporate a/biotic factors in their regression-based vital rate models (Merow et al., 2014; Ellner & Rees, 2006), they can estimate responses of populations to disturbances. Still, the fields of remote sensing, using hyperspectral and LiDAR data, and population ecology, using IPMs, have independently investigated the effect of disturbances on forests. Combining these approaches offers a way to parameterize the detail of a population model with the spatial extent of remote sensing (Rosen et al., 2025).

A third of the global above-ground carbon is stored in the boreal forests and are vulnerable to climate change (Pan et al., 2011; Palviainen et al., 2020). The increasing frequency of the historically regular (MacDonald et al., 1991; Chapin et al., 2006) wildfires is compromising the ecological integrity of the forests (Palm et al., 2022; Tollefson, 2024), even among fire-adapted tree species (Whitman et al., 2019). For example, here, we focus on black spruce, *Picea mariana* (Mill.) B.S.P., which dominates North American boreal forests. The dominance of this conifer species is mainly attributed to high post-fire replacement, as *P. mariana* has semi-serotinous cones (Fryer, 2014). However increasing fire frequency doesn’t allow as much enough time to reach reproductive maturity before the next fire, leading to reduced recruitment (Brown & Johnstone, 2012; Johnstone et al., 2020). When *P. mariana* recruitment is low, wind-dispersing species may establish instead (Johnstone et al., 2020). Successional trajectories without *P. mariana* have consequences for carbon and nutrient cycling (Melvin et al., 2015) and understory vegetation (Hart & Chen, 2006). The regeneration of this species is related to its height, as taller individuals produce more cones (Simpson & Powell, 1981). This feature makes *P. mariana* ideal to model its vital rates (*e.g*., survival, growth) as a function of individual height in IPMs (Easterling, Ellner & Dixon, 2000).

Here, we quantify the impacts of fire on the survival and growth of black spruce, *Picea mariana*, via IPMs parameterised with hyperspectral and LiDAR data. Across 26 km^2^ of Alaskan boreal forest, we examine how forest patches with different fire history inform which fire intervals are in/compatible with post-disturbance recovery. We hypothesised (H1) that lower surrounding canopy height (SCH), lower elevation, and less steep slopes would be favourable for individual survival and growth. Such conditions may support survival and growth because of the decrease in competition (Carleton & Wannamaker, 1987), temperature (Fryer, 2014), and increase in ground stability (Nicoll et al., 2005), respectively. After integrating the IPMs with the linear models chosen from testing H1, we calculated passage time and life expectancy. Specifically, we hypothesised that (H2) the longest passage time would occur in stands without recent fire history on North-facing slopes, since trees in these patches likely experience higher density dependent competition combined with limited light availability (Carleton & Wannamaker, 1987; Van Cleve et al., 1986; Nath & Ni-Meister, 2021). Also, we hypothesised (H3) that life expectancy is longest on slopes with recent fire history because the recent fire likely cleared the canopy (*P. mariana* is highly flammable; Cronan, McKenzie & Olson, 2012), replaced by new recruits. Finally, we perform a sensitivity analysis of passage time and life expectancy in relation to elevation, slope steepness, and SCH. We hypothesised (H4) that passage time and life expectancy are most sensitive to elevation where there is increased light availability at higher elevations and taller *P. mariana* individuals have been found at higher elevations on South-facing slopes (Nath & Ni-Meister, 2021).

## Materials and methods

To understand resilience of *Picea mariana* in different environmental conditions, we built IPMs parameterised with hyperspectral and LiDAR data. We used National Ecological Observatory Network data (<u>NEON Data Portal</u>) at the Caribou-Poker Creek Research Watershed (BONA site, Alaska, USA) acquired in 2017 and 2023. The overlapping area of the two scans is shown in Figure 2 B and C. Within the 26 km^2^ scan overlap, we selected areas totalling 5.5 km^2^, to include patches with different fire history, elevations and aspect (Figure 2 tiles 1-6; See supplementary materials for further details). By remotely tracking survival and growth of individuals (Section 1), we parameterised IPMs (Section 2) and estimated passage time and life expectancy (Section 3).

**Figure 1.**
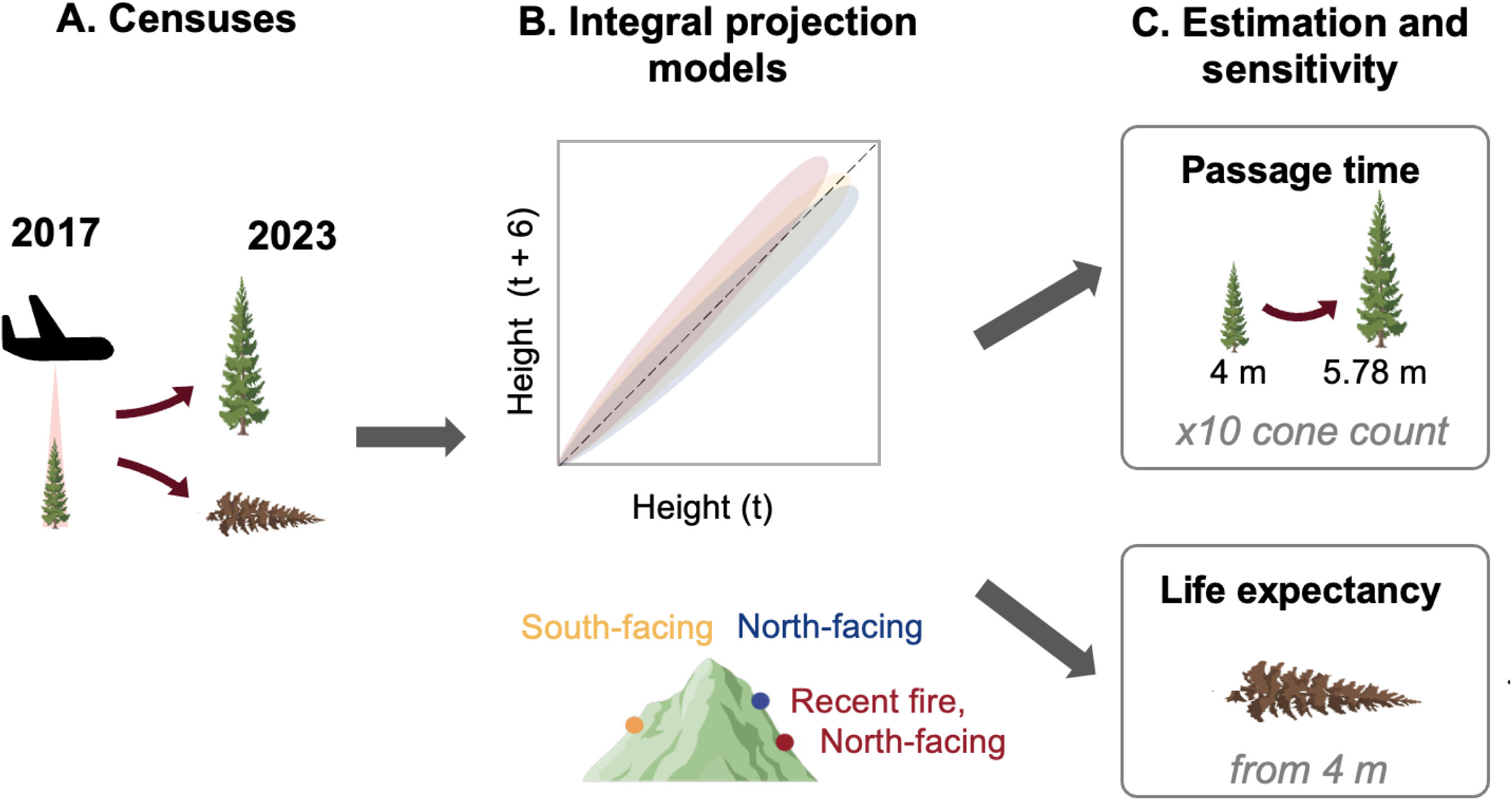
An overview of the integration of RGB, hyperspectral, and LiDAR (Light Detection and Ranging) data acquisition implemented here to estimate the time to a target height and life expectancy in a forest population of *Picea mariana*. (**A**) Here, to model the time required for boreal forests to recover from fire, we located *P. mariana* using Weinstein et al.’s (2024) species map, which used RGB imagery and hyperspectral imagery to train a machine learning model to detect individual trees in the upper canopy and identify them to species level. Next, we quantified the survival and growth of 42,816 individuals of *P. mariana* using LiDAR censuses between 2017 and 2023. (**B**) We then integrated the models of survival and growth into integral projection models (IPMs), explicitly including environmental descriptors of fire history and terrain aspects. We estimated (**C**) passage time, defined as the time to grow from 4 m to 5.78 m, which is equivalent to a 10-fold increase in cone count (Simpson & Powell, 1981), and mean life expectancy. The faster *P. mariana* can increase cone count, the more likely the stand will recover and recruit new individuals post-fire. For both passage time and life expectancy, we examined how sensitive the life history traits are to small perturbations in key environmental drivers for each of the three IPMs: elevation, slope steepness, and surround canopy height.

**Figure 2.**
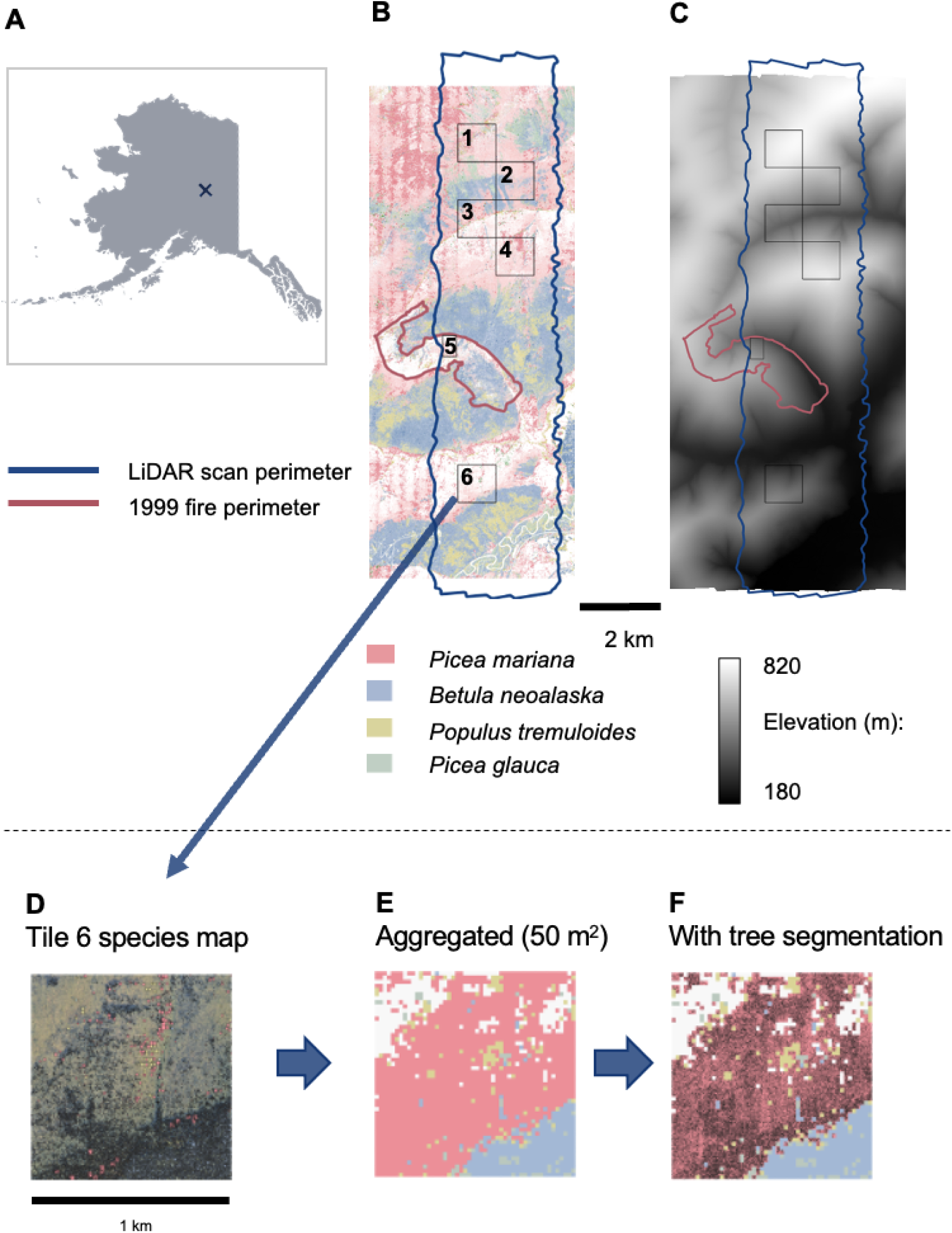
We brought together data from multiple sources, including a machine-learning based species map reliant on RGB and hyperspectral data, and satellite fire history, to locate *Picea mariana* individuals across a topographic gradient for model parameterisation. Maps of the LiDAR scan perimeter (dark blue) in 2017 and 2023 over different maps. In the (**A**) Alaska map, the LiDAR scan location is marked with an ‘X’ at the Caribou-Poker Creek Research Watershed (BONA), near Fairbanks. (**B**) The species map shows the upper-canopy species predictions (Weinstein et al., 2024) and (**C**) the elevation map shows the topography (Fischer et al., 2024) at the BONA site. The six tiles (numbered in **B**) within the scan perimeter indicate the selected areas that were examined in 2017 and 2023. Note: tile 5 was burned in 1999 and was cropped to only include trees within the fire patch (300 m x 400 m) while the other five remained 1 km x 1 km. The blue rectangle shows the overlapping LiDAR scans in 2017 and 2023. The 1999 fire perimeter (red polygon) is derived from satellite imagery (Alaskan Wildland Fire Information Map Series). Zooming in on tile 6, we illustrate the individual species labelled tree crowns over (**D**) RGB imagery based on analyses by Weinstein et al. 2024. From the species labelled tree crown map, we then (**E**) aggregated to a 50 m^2^ resolution with each grid cell labelled by species dominance, and (**F**) overlayed the LiDAR-based tree crown segmentation and filtered for trees only in *P. mariana* dominated grid cells (pink).

**Figure 3.**
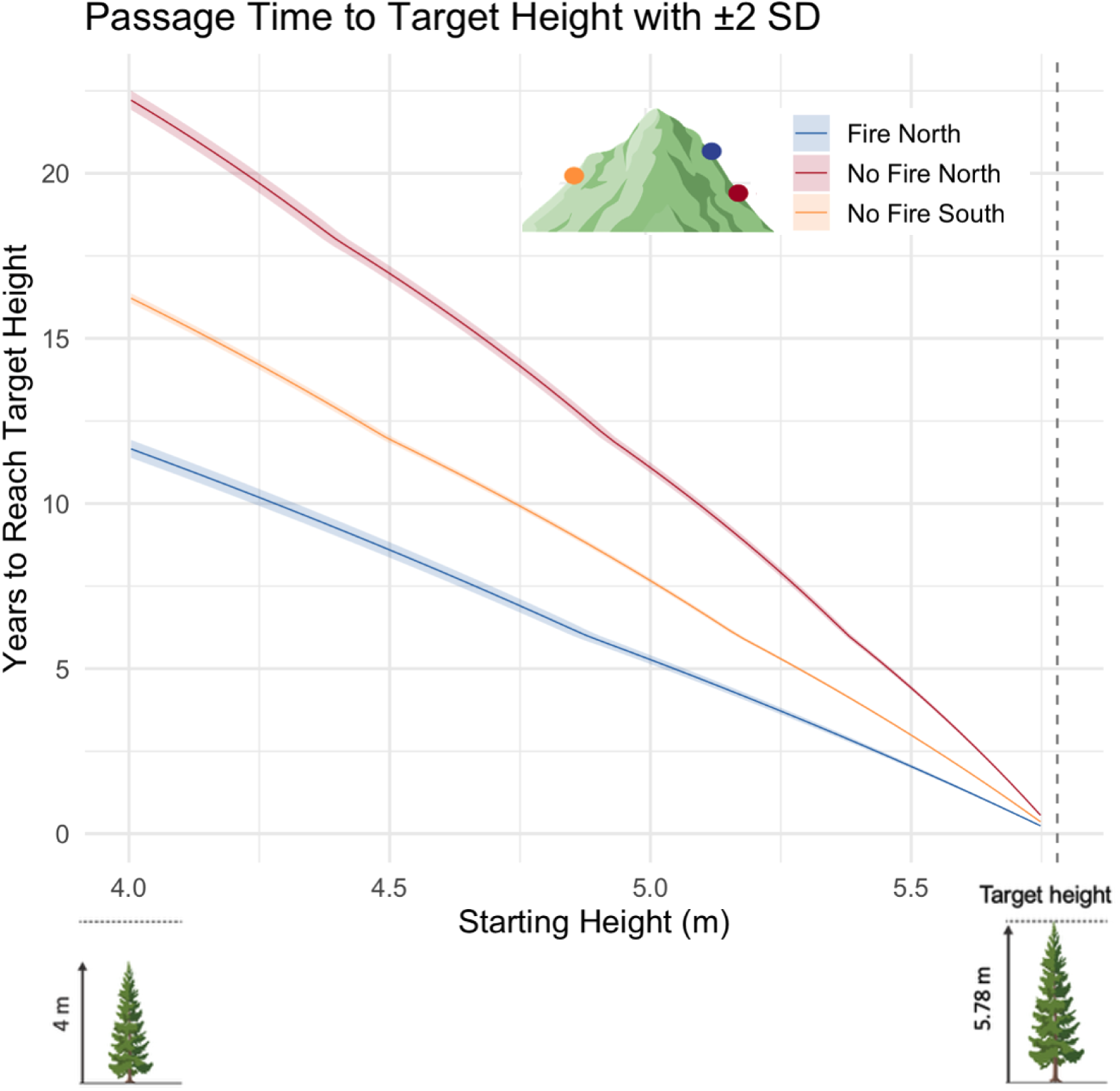
Trees on North-facing slopes with recent fire history have the fastest recovery time to heights of higher chances of self-replacement. The passage times of an individual of *Picea mariana* from 4 m tall (1.38 in log scale) to the target height of 5.78 m (1.75 in log scale; black vertical line), which informs the time required for the boreal forest to recover before fire returns. The thin shade areas correspond to 2 S.D.

### 1. Individual *P. mariana* trees

To detect individual tree crowns, we used the Dalponte and Coomes (2016) segmentation algorithm. We implemented this algorithm, which operates faster than other options (Cao et al., 2023), using the *lidR* R package (Roussel et al., 2020, 2024). Briefly, this algorithm detects high points in a canopy height model (CHM) (“local maxima”) and expands the tree crowns (“window size”) around local maxima. After detecting treetops, the algorithm segments the CHM to define the extent of the tree crown. We used a 1 m resolution CHM modelled with the “highest” algorithm in Fischer et al. (2024), which is designed to reveal treetops in the upper canopy. We only tracked the trees with a crown area of at least 3 m^2^ because those trees heights would be informed by at least four CHM pixels. Individuals with a smaller crown area have high height variability with any misalignment of census scans. Please see Section 1 of the Supplementary Materials for full details about of the algorithm fine tuning (24.4% of auto-segmented trees with a crown area of at least 3 m^2^ overlapped with manually segmented trees).

To select for *P. mariana*, we filtered out areas dominated by other species within each of the six examined tiles. The upper canopy of the BONA site includes three other species: Alaskan birch (*Betula neoalaskana*), trembling aspen (*Populus tremuloides*), and white spruce (*Picea glauca*). Weinstein et al.’s (2024) hyperspectral species map allowed us to differentiate patches dominated by *P. mariana* from patches dominated by other species. Briefly, Weinstein and colleagues used 1 m resolution hyperspectral reflectance and geolocated field data to build a machine learning model that assigned species-IDs to tree crowns segmented with 1 m resolution RGB imagery (Weinstein et al., 2020, 2024). The BONA site had an 82% “micro-accuracy” (indication of performance in labelling the more common species), which is the third highest site score out of 23 NEON sites (see Table 2 of

Weinstein et al. 2024). To locate non-*P. mariana* upper canopy trees, we created an aggregated version of species map. We created a 50 m^2^ raster layer that labelled raster cells based on the most common species, which further ensured removal of non-*P. mariana* tree crowns (Figure 2E). We obtained a dataset of 42,816 individuals with a crown area over 3 m^2^ (Figure 2F) within the *P. mariana-*dominated 50 m^2^ cells. The average crown area was 8.01 m^2^, within the range observed in the NEON field data for this species (NEON, 2024). See Section 2 in the Supplementary Materials for full details about the tile selection criteria.

To test our hypotheses, we carried out spatial analyses and statistical modelling on R version 4.3.2 (R Core Team, 2023). We implemented the spatial data processing using the *sf* (Pebesma, 2018) and *terra* R packages (Hijmans et al., 2024) for vector (tree crowns) and raster data (CHMs and DTMs), respectively.

### 2. LiDAR-IPM framework

#### 2.1. Model parameters

##### Survival and growth

To estimate survival and growth, we measured height change between 2017 and 2023. Tree height was the maximum CHM height within each tree polygon. We used the CHM calculated from the “spikefreel” algorithm, which supports robust comparisons between acquisitions (Fischer et al., 2024). In the absence of fire, the main causes of mortality are windthrow and waterlogging from sinking in to melted permafrost (Dearborn et al., 2021; Elie & Ruel, 2005). We classified an individual as dead if its height decreased by ≥50% between 2017 and 2023. While past studies have used a 30% threshold, we were more conservative to match the high variability in height measurements between censuses (Battison et al., 2024 (30%); Ma et al., 2023 (30-50%); Duncanson & Dubayah, 2018 (30%)). See Section 3 of the Supplementary Materials for full details about the height change and mortality rate observed.

##### Environmental drivers

We quantified each environmental driver to investigate (H1-3) their effect on survival and growth. Recent fire was a binary factor, with “no” for individuals located outside of the 1999 fire patch (the last fire occurred before 1940; Bureau of Land Management, n.d.). For aspect and slope, we transformed the 2017 Digital Terrain Model (DTM) into an aspect and slope map using the R *terra* package (Hijmans et al., 2024). For aspect, we then cosine-transformed the map from a 360° scale to −1 to 1 scale to focus on North and South aspect, respectively (Beese et al., 2022). We extracted the mean elevation for each tree from the 2017 DTM. Lastly, we measured the Surround Canopy Height (SCH; Krause & Lemay, 2023; Battison et al., 2024). SCH here is defined as the average height within a 15 m buffer around each tree. Our findings below are insensitive to the choice of SCH radius, which is shown in Section 4 of the Supplementary Materials.

To also consider if height changes may be due to the differences between the points density of the scans in 2017 and 2023, we determined the Relative Change in Point density (RCP, Eq. 1) for each tree. The average point density in 2023 was ∼four-fold higher than in 2017 (20.1 pts/m^2^ in 2023, 5.12/m^2^ in 2017; <u>NEON data portal</u>). We used a relative metric because higher point densities do plateau to a stable height measurement (Hawryło et al. 2024).

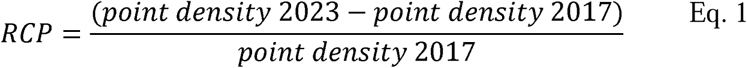

#### 2.2. Vital rate models

To estimate passage time and life expectancy, we parameterised integral projection models (IPMs; Easterling, Ellner & Dixon, 2000). IPMs are demographic models that describe changes in the structure of a population along a continuous trait (here, tree height) in discrete time. The integration of survival and growth models into the IPM framework produces a two-dimensional *P* kernel (Ellner, Childs & Rees, 2016). IPMs often contain an *F* kernel, which describes the recruitment of new individuals to the population (Easterling, Ellner & Dixon, 2000; Ellner & Rees, 2006). However, only survival and growth are necessary to test our hypotheses (but see Discussion; Merow et al., 2014; Metcalf et al., 2013). Our *P* kernel tracks progression of the structure of a *P. mariana* population, *n*, between a range of possible heights, from the lower (*L*) to upper (*U*) bounds of the integral defined by Equation 2, to the next time step in the censused data (here *t* + 6 years):

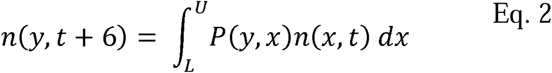

Equation 2 can be broken down into equations 3-5 to obtain the expected distribution of tree heights at time *t*+ 6 for an individual of height *x*,at time *t*, conditional on having survived. For more details about IPM midpoint integration, including determination of the lower (*L*) and upper (*U*) bounds of the *P* kernel integral (Eq. 2) and choice of meshpoints, see Section 5 of the Supplementary Materials.

The *P* kernel is composed of the *s* survival function (probability of survival for a given height *x* in time *t*) and the *g* growth function (the probability that a tree of a given height *x* in time *t* will grow to a given height *y* in *t*+6):

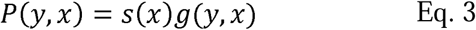

To build the components of the *P* kernel, we selected the most parsimonious models of survival and growth. A description of all environmental covariates derived from remote sensing data is found in Table S1. Before model construction, we checked for multicollinearity in the model predictors. However, we found no multicollinearity high enough to justify removing any predictor from our models (See Figure S5).

To test H1 on the effect of environmental drivers on survival and growth and to select the models for the IPMs (H2-4), we started with the most complex additive models given our hypotheses. We modelled the probability that a tree survived in the examined period as a binomial logistic regression function of height *x*, in log scale. (Eq. 4) and we modelled growth as the change in height of surviving individuals (Eq. 5). The final models were polynomial (logistic, in the case of survival) regressions, including a quadratic term for height, a change in intercept for individuals on North-facing slopes (*North*) or individuals in an area with a recent fire (*Fire*; inside a stand burnt in 1999) and four environmental covariates (*Elevation*, *Steepness, RCP,* and *SCH*). The four environmental covariate terms are the mean value of that environmental variable across the dataset (*e.g. Elevation,* See Section 8 of the Supplementary Materials) multiplied by the estimate/slope of that environmental variable, indicated by the hat (*e.g.*ê_S_). Full summaries of selection process and model estimates can be found in the Supplementary Material (Sections 9-11). Finally, we quantified the error in the mean growth function, to capture the magnitude of uncertainty in LiDAR acquisitions (Clark, 2003), which is further detailed in Section 11 of the Supplementary Materials. We conducted ANOVA analyses and results of the F-test for the environmental drivers of growth are reported with support from the library “report” (Makowski et al. 2025).

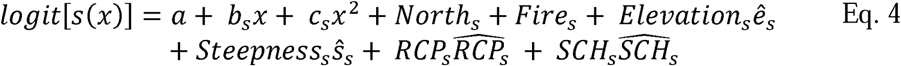

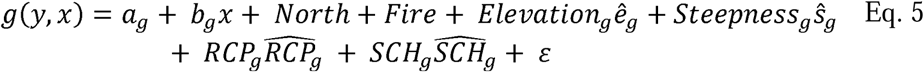

To test hypotheses regarding the difference between estimations of passage time (H2) and mean life expectancy (H3), we altered the recent fire history and aspect conditions between IPMs. The resulting three IPMs were two North-facing (1) with or (2) without recent fire history, and one additional IPM with (3) South-facing without recent fire history. We did not parameterise an IPM with recent fire and South-facing slope, as this combination of conditions does not exist at the BONA site during the monitored period. The same survival and growth functions were used to build all three IPMs (Eq. 4-5 integrated as in Eq. 3) but differed in intercepts for fire history and aspect. The final integrated IPM kernels are also presented in the Supplementary Material in Section 12. Because of the six-year interval between censuses, the outputs of passage time and mean life expectancy were multiplied by six, so the final outputs are in the units of years.

### 3. Passage time and life expectancy

To test our hypothesis that (H2) the longest passage time would be located on North-facing slopes without recent fire history, we compared the respective IPM passage times, that is, the time elapsed for individuals of a given height to achieve a target height (Merow et al., 2014). Our passage time of interest was between 4 m, corresponding to the shortest individual tree detected in our dataset, to a target height of 5.78 m tall. The chosen target height corresponds to a 10-fold increase in cone production compared to a 4 m tall individual tree, according to the height to cone-count regression modelled by Simpson & Powell (1981). With more cones, taller trees may have a higher chance at self-replacement (Simpson & Powell, 1981; Viglas, Brown & Johnstone, 2013). Exploring passage time to a height of 5.78 m also prevents the model from extrapolating outside of the maximum height observed in the single fire patch (See Section 13 of the Supplementary Materials). To estimate passage time, we started with a population vector with 1000 individuals in the cell representing the starting height. We then applied the *P* kernel iteratively to track the number of transitions for the mean of the population distribution to reach the target height, allowing for linear interpolation. We performed 1000 bootstraps, sampling 95% of the data each time to get the standard deviation. To test the hypothesis (H3) that life expectancy is highest on slopes with no recent fire history, regardless of aspect, we used the *Rage R* package (Barks et al., 2023; Jones et al., 2022). Here, life expectancy is defined as the mean age at death for individuals, conditional on having reached 4 m tall, the shortest height observed with LiDAR in this study. Since fire did not occur between the censuses, these are estimates of life expectancy in the absence of fire. To understand the variance of mean life expectancy between environmental conditions, we bootstrapped 95% of the individual trajectories a hundred times. We then calculated standard errors from these bootstraps to detect significant differences in mean life expectancy. To test whether the mean life expectancy was significantly different between conditions, we performed a one-way ANOVA followed by a post-hoc HSD Tukey test.

For both life expectancy and passage time, we carried out a sensitivity analysis to test (H4) that both life history traits are most sensitive to elevation, with higher sensitivity on North-facing slopes. For each of the three IPMs, the three environmental drivers (elevation, slope steepness, and surround canopy height) were perturbed using the ‘brute force’ method (*sensu* Morris & Doak, 2002), by increasing by the coefficient (slope)t of each parameter by 1% one at a time to then obtain a new estimate of passage time and life expectancy. The impact of each perturbation was determined by comparing the new estimate to the original estimates of mean life expectancy and passage time.

## Results

### 1. Environmental drivers of survival and growth

We found partial support for our hypothesis (H1) that certain environmental conditions would positively influence the survival and growth of individuals of *P*. *mariana* (Table 1). Specifically, lower slope steepness (*i.e.*, flatter terrain) has a positive effect on both survival (OR = 0.82, 95% CI = [0.730, 0.920]) and growth (F_1, 42,463_ = 141.41, p < 0.001; Eta2 (partial) = 0.0003, [0.0002, 1.000]). Contrary to our hypothesis, though, increases in elevation and surround canopy height have a positive effect on both survival (Elevation: (OR = 0.82, 95% CI = [0.730, 0.920]; SCH: OR = 4.68, 95% CI = [3.790, 5.840]) and growth (Elevation: (F_1,42463_ = 11.68, p < 0.001; Eta2 (partial) < 0.001, [0.000, 1.000]); SCH: (F_1, 42463_ = 3823.95, p < 0.001; Eta2 (partial) = 0.08, [0.080, 1.000])). It is worth noting that, in both models of survival and growth, the slope estimates are quite small for the overall regression slope, but that their significant differences and the AIC comparison support their importance to the model in improving population forecasts (Tables S2-8). Full Odds Ratio and ANOVA analyses for all parameters of the survival and growth models are available in the Supplementary Online Materials in Sections 8 and 9, respectively.

**Table 1.**
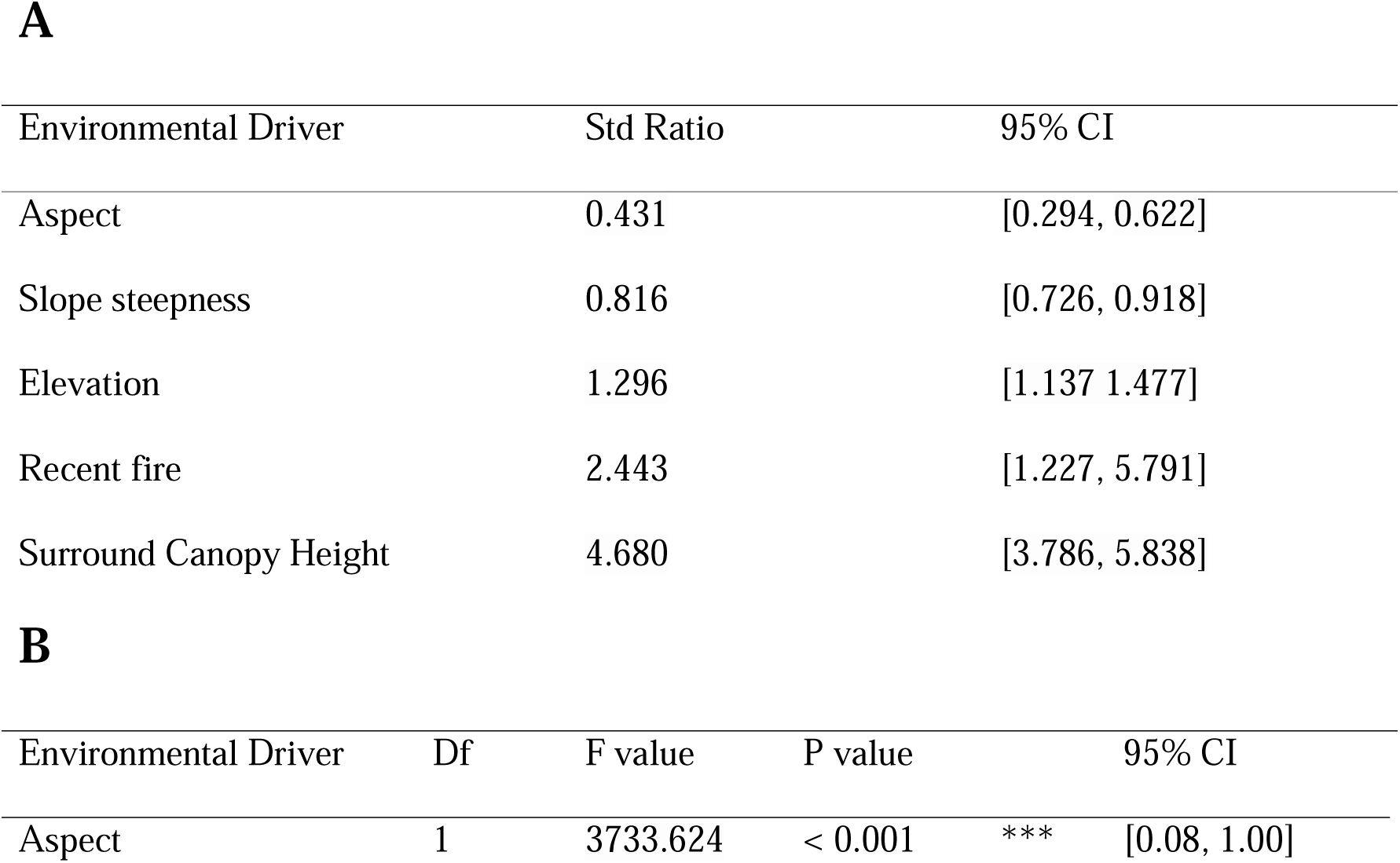
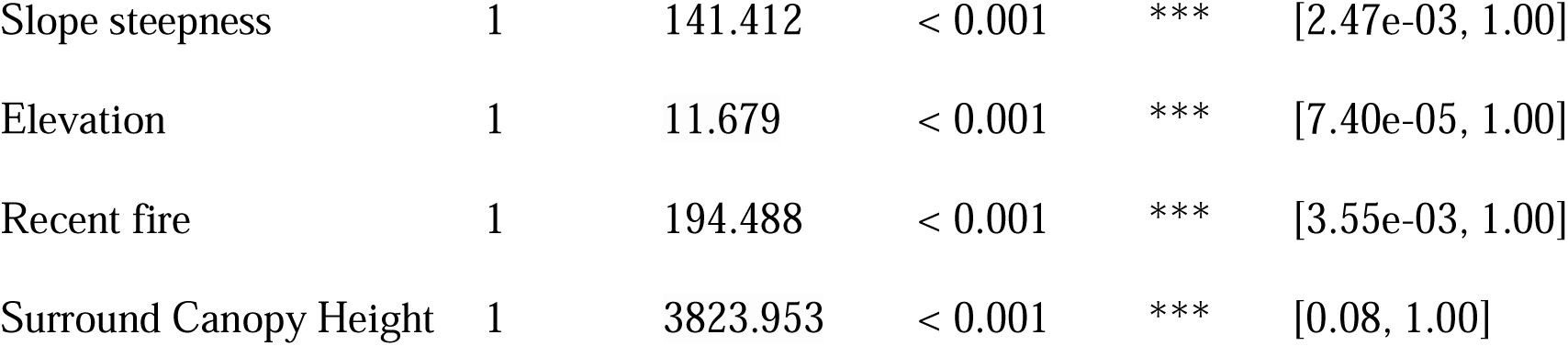
All five environmental drivers examined in this study predict the survival of *Picea mariana* individuals between 2017 and 2023, as shown by Odds Ratio calculations (A) and by ANOVA analysis (B). *** P < 0.001. We calculated the environmental drivers in the same way for both models: elevation is the high above sea level (m) which we derived from the 2017 digital terrain model (DTM); slope steepness is the percentage change in the DTM, aspect is a binary variable of either North- or South-facing, again derived from the 2017 DTM; surround canopy height (SCH) is the average height of the canopy within a 15m radius of the tree, which we derived from the 2017 canopy height model (CHM); and recent fire history was assigned to trees growing inside the 1999 fire patch, as provided by an Alaska Wildland Fire Information Map Series shapefile. Full model summaries including all model parameters and associated analyses can be found in the Supplementary Online Materials in Sections 9-11.

### 2. Passage time and sensitivity analysis

As we had expected (H2), the longest passage time occurs in stands without recent fire history on North-facing slopes. Specifically, stands located on North-facing slopes without recent fire history have a passage time of 22.21 years (± 0.15 SD). The passage times in other conditions are lower with a passage time of 16.22 years (± 0.14) on South-facing slopes without recent fire history and 11.65 years (± 0.08) for sites with recent fire history on a North-facing slope.

We did not find support for our hypothesis (H4) that passage time would be most sensitive to elevation. Indeed, across all combinations of environmental conditions, increases in the elevation slope parameter correspond to short passage time, as mediated by its effects on survival and growth (Figure 5A-C). However, passage is most negatively sensitive to surround canopy height, as mediated by its effect on growth (Figure 5B). The negative sensitivity passage time to perturbations across all environmental drivers is most extreme for the no recent fire and North-facing combination, followed by no recent fire, South facing and the least sensitivity in the recent fire and North-facing conditions.

### 3. Life expectancy and sensitivity analysis

Contrary to our third hypothesis (H3), life expectancy is estimated to be much longer in stands located on slopes without recent fire history, rather than with recent fire history. The conditions of recent fire on a North-facing slope result in a mean life expectancy of 579.37 years (± 5.30 S.E.), while no recent fire history stands on North- and South-facing slopes raise a mean life expectancy of 1,198.27 years (± 4.24) and 1,549.02 years (± 5.72), respectively (Figure 4). The one-way ANOVA confirms significant difference in the mean life expectancies between at least two of the groups (*F*_2, 297_ = 9,177.23, *P* < 0.001; 95% CI [0.980, 1.000]). Furthermore, in the post-hoc Tukey test, all possible comparisons between two groups are significantly different (adjusted *P*-values for each comparison < 0.050).

**Figure 4.**
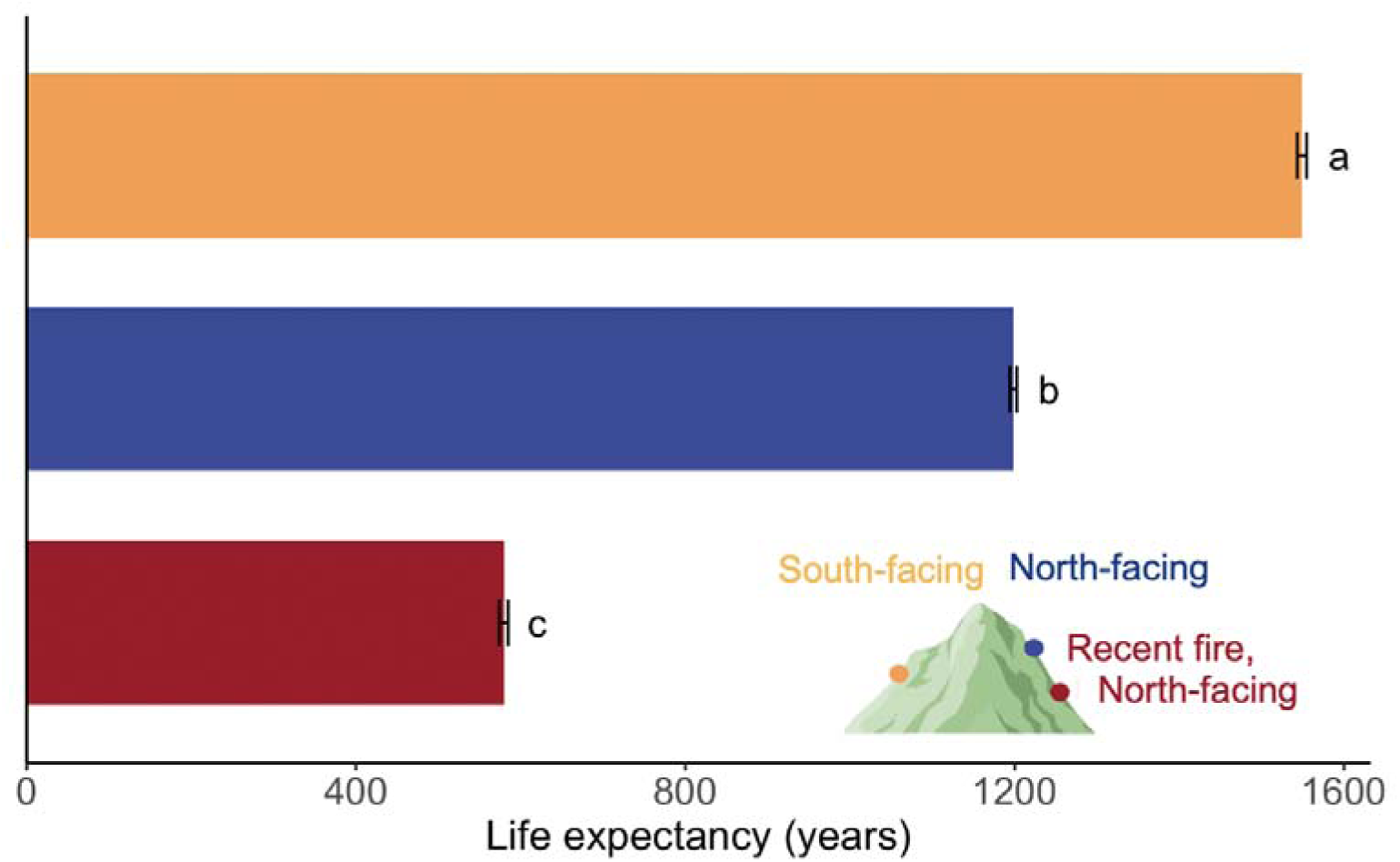
Life expectancy of *Picea mariana* varies significantly across patches exposed to different aspects (South *vs.* North) and fire history. Specifically, in more recently burned stands, individuals do not live as long as those in a not recently burned stand of the same height. Letters correspond to a post-hoc Tukey test, where groups not sharing the same letter are statistically significantly different at *P* < 0.05.

**Figure 5.**
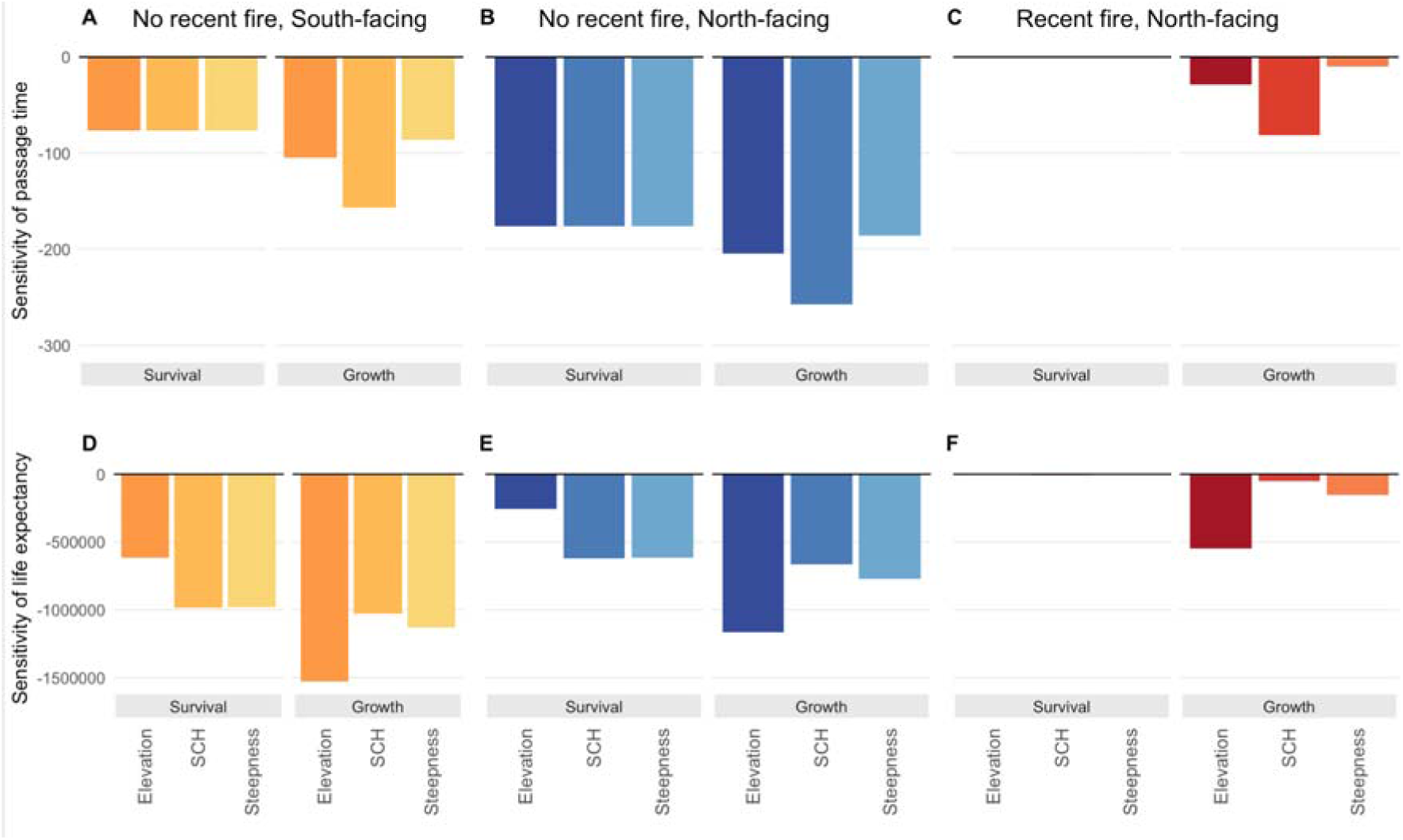
Passage time and life expectancy of *Picea mariana* under the three considered environmental conditions vary in sensitivity to the three environmental drivers: elevation, surround canopy height (SCH), and steepness, in both the vital rates of survival and growth. The environmental conditions modelled are: No recent fire, South-facing (**A** and **D**); No recent fire, North-facing (**B** and **E**); Recent fire, North-facing (**C** and **F**). All three environmental conditions have a negative sensitivity to all environmental drivers via contributions of both survival and growth for both passage time and life expectancy. The environmental conditions with the highest sensitivity to any perturbation in environmental drivers for passage time is No recent fire on a North-facing slope (Panel **B**), while the environmental conditions with the highest sensitivity to any perturbation in environmental drivers for life expectancy is no recent fire on a South-facing slope (Panel **D**).

We found support for our fourth hypothesis (H4) that life expectancy should be highly sensitive to elevation. Indeed, elevation was the primary driver of life expectancy via its effects on growth across all environmental conditions (Figure 5 D-F). The negative sensitivity life expectancy to perturbations across all environmental drivers is most extreme for the no recent fire and South-facing combination, followed by no recent fire, North-facing and the least sensitivity in the recent fire and North-facing conditions.

## Discussion

Anthropogenic disturbances, including pests, drought, and fires, are threatening tree longevity (Cobb et al., 2017) and consequently forest productivity globally (Morin et al., 2018; Hogan et al., 2024). To maintain forest biodiversity and its multiple ecosystem services (Mo et al., 2023; Brockerhoff et al., 2017), we urgently need to identify which forests are least resilient – and to what disturbances – so we can develop cost-effective, tailor-made management solutions (Thorogood et al., 2023; Ibáñez et al., 2019). However, the survival and growth rates of trees can vary greatly among and within species (Bialic-Murphy et al., 2024; Drobyshev et al., 2013) and across environmental conditions (Ohse et al., 2023; McMahon, Parker & Miller, 2010). To overcome this challenge, here we parameterised a LiDAR-based integral projection model (IPM; Easterling, Ellner & Dixon, 2000) to examine the drivers of passage time and life expectancy of *Picea mariana*, a species vulnerable to increasing fire frequency and severity (Brown & Johnstone, 2012; Johnstone et al., 2020). We found that the combination of conditions that support the fastest growth (and thus shortest passage times) also incur the shortest life expectancy. Passage time is most sensitive to perturbations in surround canopy height, while life expectancy is most sensitive to perturbations in elevation.

The stands modelled on North-facing slopes recently affected by fire produced the shortest passage time and life expectancy. Such conditions may expect to increase their cone production ten-fold (*i.e.*, from 4 m to 5.8 m height; Simpson and Powell, 1987) in less time than stands without recent fire history. This aligns with the finding that a site burned in 2004 in interior Alaska, stands of *P. mariana* grew faster on burned sites compared to unburned sites (Tsuyuzaki et al., 2014). While we had hypothesised that such conditions would also support longer life expectancy, we found the opposite. Though unexpected, the finding aligns with the general life history pattern of the fast-slow trade off, which is well documented in trees (Rüger et al., 2020), and more broadly the Plant Kingdom (Salguero-Gómez et al., 2016). Possibly, slower growth can support higher wood density and the mechanical strength to resist breakage/mortality by snowfall or wind exposure at higher elevations (Elie & Ruel, 2005; Chave et al., 2009). Stands with higher basal area and more mature stands were found to have greater viable seed rain after fire in the Northwest Territories Canada (Reid et al., 2023), so understanding how the conditions that support life expectancy in a trade-off with passage time is important. Our estimates of relatively long life expectancies in stands not exposed to recent fires are consistent with estimates of over 2,000 years for stands of this species in Quebec, Canada, which have survived previous fire exposure (Nappi et al., 2010).

Incorporating recruitment data into our LiDAR-parameterised IPMs will further inform our understanding of long-term indicators of resilience of these diverse forests. We would need to estimate reproduction of established individuals as well as the survival and growth of seedlings to the threshold of 4 m height, where LiDAR products can detect individual tree height and canopy area (Dalponte & Coomes, 2016; Cao et al., 2023). Importantly though, being taller is not required for black spruce to produce cones, though taller individuals bare more cones (Simpson & Powell, 1981). While a wealth of *P. mariana* recruitment data exists (Baltzer et al., 2021; Dearborn et al., 2021), the larger challenge remains to ‘close the life cycle’ with a complete picture of environmental drivers at play (Lloyd et al., 2005). Furthermore, future field work would allow us to partition the biological variation from the CHM or segmentation error that create uncertainty in passage time and life expectancy estimations. The integration of multi-sensed, multi-platform data acquisition techniques and pertinent *in situ* validation paint an exciting future at the interface of novel technologies and ecological monitoring in forests (Qi et al., 2025; Rosen et al., 2025).

We anticipate that IPMs that include complete data of the full life cycle of a target species, its environmental drivers, and community-level interactions, as done recently (Gupta et al., 2024), will lead the way in the calculation of key metrics of demographic resilience to pulse disturbances, like fire. These interdisciplinary integrations could be more generally applied to other ecosystems such as coral reefs or tundra landscapes (Capdevila et al., 2020; Cant et al., 2023). These integrative approaches will also allow us to spatially map these demographic processes out onto a landscape as modelled to locate stands that are ecologically vulnerable (Merow et al., 2014; Viereck et al., 2008). In bringing together demographic and environmental data, including LiDAR-derived estimates for larger individual trees and tiny seedlings, we will be able to advance our understanding of global trends, where boreal systems are currently underrepresented (Rotbarth et al., 2025; Reeb, 2024; Ohse et al., 2023). The resilience of forests to Anthropogenic disturbances depends on the environmental conditions in which its populations have become adapted to survive and grow to maturity (Ibáñez et al., 2019). Our analyses reveal that the life history traits that aid pulse disturbance recovery, including shorter passage time to heights of higher reproductive viability and longer life expectancy, are further mediated by elevation, SCH and slope steepens. Our analytical pipeline, which integrates hyperspectral-species maps and LiDAR-derived heights of individual trees, increases the power of the population dynamics models to be observed across combinations of environmental conditions. We encourage the research community to apply our pipeline to understand other environmental threats where appropriate remote sensing data are available.

## Supporting information

SOM

## Acknowledgements

We thank R. Battison for LiDAR segmentation and height tracking code, and B. Weinstein and colleagues and F. Fischer for sharing data products of species crown maps and digital surface models respectively. J.M. was supported in part by the Mike Soper Bursary Fund from the Department of Biology at the University of Oxford. We thank the members of the SalGo team and Selva lab for their input in early stages of development of this project. AR was supported by the Oxford BBSRC DTP studentship (BB/T008784/1) in RSG’s lab. RSG was supported by a NERC Pushing the Frontiers grant (NE/X013766/1).

## Supporting Information

“*Remote Sensing in Ecology and Conservation* expects that data supporting the results in the paper will be archived in an appropriate public repository.”

## Authors’ Contribution Statement

JMM, RSG, and TJ conceived the ideas and designed methodology; JMM sourced the data and analyzed the data; RSG produced R scripts for the demographic modelling and AR and JMM adapted them; JMM led first draft of the manuscript under the guidance of RSG; All authors contributed critically to the drafts and gave final approval for publication.

## References

Baltzer, J.L., Day, N.J., Walker, X.J., Greene, D., Mack, M.C., et al. (2021) Increasing fire and the decline of fire adapted black spruce in the boreal forest. Proceedings of the National Academy of Sciences. 118 (45), e2024872118. doi:10.1073/pnas.2024872118.

Barks, P., Buss, D., Capdevila, P., Caswell, H., Che-Castaldo, J.P., Hinrichsen, R.A., Jackson, J., James, T., Jones [aut, O., cre, Levin, S., Petry, W.K., Salguero-Gomez, R., Schuette, C., Stott, I., Thomas, C.C. & Hernández, C.M. (2023) Rage: Life History Metrics from Matrix Population Models. https://cran.r-project.org/web/packages/Rage/index.html.

Battison, R., Prober, S.M., Zdunic, K., Jackson, T.D., Fischer, F.J. & Jucker, T. (2024) Tracking tree demography and forest dynamics at scale using remote sensing.p.2024.06.11.598435. doi:10.1101/2024.06.11.598435.

Beese, L., Dalponte, M., Asner, G.P., Coomes, D.A. & Jucker, T. (2022) Using repeat airborne LiDAR to map the growth of individual oil palms in Malaysian Borneo during the 2015–16 El Niño. International Journal of Applied Earth Observation and Geoinformation. 115, 103117. doi:10.1016/j.jag.2022.103117.

Bialic-Murphy, L., McElderry, R.M., Esquivel-Muelbert, A., van den Hoogen, J., Zuidema, P.A., et al. (2024) The pace of life for forest trees. Science. 386 (6717), 92–98. doi:10.1126/science.adk9616.

Brockerhoff, E.G., Barbaro, L., Castagneyrol, B., Forrester, D.I., Gardiner, B., González-Olabarria, J.R., Lyver, P.O., Meurisse, N., Oxbrough, A., Taki, H., Thompson, I.D., van der Plas, F. & Jactel, H. (2017) Forest biodiversity, ecosystem functioning and the provision of ecosystem services. Biodiversity and Conservation. 26 (13), 3005–3035. doi:10.1007/s10531-017-1453-2.

Brown, C.D. & Johnstone, J.F. (2012) Once burned, twice shy: Repeat fires reduce seed availability and alter substrate constraints on *Picea mariana* regeneration. Forest Ecology and Management. 266, 34–41. doi:10.1016/j.foreco.2011.11.006.

Canelles, Q., Aquilué, N., James, P.M.A., Lawler, J. & Brotons, L. (2021) Global review on interactions between insect pests and other forest disturbances. Landscape Ecology. 36 (4), 945–972. doi:10.1007/s10980-021-01209-7.

Cant, J., Capdevila, P., Beger, M. & Salguero-Gómez, R. (2023) Recent exposure to environmental stochasticity does not determine the demographic resilience of natural populations. Ecology Letters. 26 (7), 1186–1199. doi:10.1111/ele.14234.

Cao, Y., Ball, J.G.C., Coomes, D.A., Steinmeier, L., Knapp, N., Wilkes, P., Disney, M., Calders, K., Burt, A., Lin, Y. & Jackson, T.D. (2023) Benchmarking airborne laser scanning tree segmentation algorithms in broadleaf forests shows high accuracy only for canopy trees. International Journal of Applied Earth Observation and Geoinformation. 123, 103490. doi:10.1016/j.jag.2023.103490.

Capdevila, P., Stott, I., Beger, M. & Salguero-Gómez, R. (2020) Towards a Comparative Framework of Demographic Resilience. Trends in Ecology & Evolution. 35 (9), 776–786. doi:10.1016/j.tree.2020.05.001.

Carleton, T.J. & Wannamaker, B.A. (1987) Mortality and Self-thinning in Postfire Black Spruce. Annals of Botany. 59 (6), 621–628. doi:10.1093/oxfordjournals.aob.a087358.

Cavendar-Bares, J., Gamon, J.A. & Townsend, P.A. (2020) Remote Sensing of Plant Biodiversity. Springer Nature. https://library.oapen.org/handle/20.500.12657/39986.

Chapin, F.S., Oswood, M.W., Van Cleve, K., Viereck, L.A. & Verbyla, D.L. (2006) Alaska’s Changing Boreal Forest. Oxford University Press. doi:10.1093/oso/9780195154313.001.0001.

Chave, J., Coomes, D., Jansen, S., Lewis, S.L., Swenson, N.G. & Zanne, A.E. (2009) Towards a worldwide wood economics spectrum. Ecology Letters. 12 (4), 351–366. doi:10.1111/j.1461-0248.2009.01285.x.

Choi, D.H., LaRue, E.A., Atkins, J.W., Foster, J.R., Matthes, J.H., Fahey, R.T., Thapa, B., Fei, S. & Hardiman, B.S. (2023) Short-term effects of moderate severity disturbances on forest canopy structure. Journal of Ecology. 111 (9), 1866–1881. doi:10.1111/1365-2745.14145.

Clark, J.S. (2003) Uncertainty and Variability in Demography and Population Growth: A Hierarchical Approach. Ecology. 84 (6), 1370–1381.

Cobb, R.C., Ruthrof, K.X., Breshears, D.D., Lloret, F., Aakala, T., et al. (2017) Ecosystem dynamics and management after forest die-off: a global synthesis with conceptual state-and-transition models. Ecosphere. 8 (12), e02034. doi:10.1002/ecs2.2034.

Cronan, J., McKenzie, D. & Olson, D. (2012) *Fire regimes of the Alaskan boreal forest*.

Cushman, K.C., Burley, J.T., Imbach, B., Saatchi, S.S., Silva, C.E., Vargas, O., Zgraggen, C. & Kellner, J.R. (2021) Impact of a tropical forest blowdown on aboveground carbon balance. Scientific Reports. 11 (1), 11279. doi:10.1038/s41598-021-90576-x.

Dalponte, M. & Coomes, D.A. (2016) Tree-centric mapping of forest carbon density from airborne laser scanning and hyperspectral data. Methods in Ecology and Evolution. 7 (10), 1236–1245. doi:10.1111/2041-210X.12575.

Dearborn, K.D., Wallace, C.A., Patankar, R. & Baltzer, J.L. (2021) Permafrost thaw in boreal peatlands is rapidly altering forest community composition. Journal of Ecology. 109 (3), 1452–1467. doi:10.1111/1365-2745.13569.

Drobyshev, I., Gewehr, S., Berninger, F. & Bergeron, Y. (2013) Species specific growth responses of black spruce and trembling aspen may enhance resilience of boreal forest to climate change. Journal of Ecology. 101 (1), 231–242. doi:10.1111/1365-2745.12007.

Duncanson, L. & Dubayah, R. (2018) Monitoring individual tree-based change with airborne lidar. Ecology and Evolution. 8 (10), 5079–5089. doi:10.1002/ece3.4075.

Easterling, M., Ellner, S. & Dixon, P. (2000) Size-Specific Sensitivity: Applying a New Structured Population Model. Ecology. 81, 694–708. doi:10.2307/177370.

Elie, J.-G. & Ruel, J.-C. (2005) Windthrow hazard modelling in boreal forests of black spruce and jack pine. Canadian Journal of Forest Research. 35 (11), 2655–2663. doi:10.1139/x05-189.

Ellner, S.P., Childs, D.Z. & Rees, M. (2016) *Data-driven Modelling of Structured Populations: A Practical Guide to the Integral Projection Model*. Lecture Notes on Mathematical Modelling in the Life Sciences. Cham, Springer International Publishing. doi:10.1007/978-3-319-28893-2.

Ellner, S.P. & Rees, M. (2006) Integral projection models for species with complex demography. The American Naturalist. 167 (3), 410–428. doi:10.1086/499438.

Fassnacht, F.E., White, J.C., Wulder, M.A. & Næsset, E. (2024) Remote sensing in forestry: current challenges, considerations and directions. Forestry: An International Journal of Forest Research. 97 (1), 11–37. doi:10.1093/forestry/cpad024.

Fischer, F.J., Jackson, T., Vincent, G. & Jucker, T. (2024) Robust characterisation of forest structure from airborne laser scanning—A systematic assessment and sample workflow for ecologists. Methods in Ecology and Evolution. n/a (n/a). doi:10.1111/2041-210X.14416.

Fryer, J. (2014) Fire regimes of Alaskan black spruce communities. Fire Effects Information System; US Department of Agriculture, Forest Service, Rocky Mountain Research Station, Fire Sciences Laboratory (Producer).

Gupta, A., Gascoigne, S., Barabás, G., Qi, M., Fenollosa, E., Thornley, R., Hernandez, C., Hector, A. & Salguero-Gómez, R. (2024) Variation in precipitation drives differences in interactions and short-term transient instability between grassland functional groups: a stage-structured community approach.p.2024.10.07.617067. doi:10.1101/2024.10.07.617067.

Hammond, W.M., Williams, A.P., Abatzoglou, J.T., Adams, H.D., Klein, T., López, R., Sáenz-Romero, C., Hartmann, H., Breshears, D.D. & Allen, C.D. (2022) Global field observations of tree die-off reveal hotter-drought fingerprint for Earth’s forests. Nature Communications. 13 (1), 1761. doi:10.1038/s41467-022-29289-2.

Hart, S.A. & Chen, H.Y.H. (2006) Understory Vegetation Dynamics of North American Boreal Forests. Critical Reviews in Plant Sciences. 25 (4), 381–397. doi:10.1080/07352680600819286.

Hijmans, R.J., Bivand, R., Dyba, K., Pebesma, E. & Sumner, M.D. (2024) terra: Spatial Data Analysis. https://cran.r-project.org/web/packages/terra/index.html.

Hogan, J.A., Domke, G.M., Zhu, K., Johnson, D.J. & Lichstein, J.W. (2024) Climate change determines the sign of productivity trends in US forests. Proceedings of the National Academy of Sciences. 121 (4), e2311132121. doi:10.1073/pnas.2311132121.

Ibáñez, I., Acharya, K., Juno, E., Karounos, C., Lee, B.R., McCollum, C., Schaffer-Morrison, S. & Tourville, J. (2019) Forest resilience under global environmental change: Do we have the information we need? A systematic review. PLOS ONE. 14 (9), e0222207. doi:10.1371/journal.pone.0222207.

Johnstone, J.F., Celis, G., Chapin III, F.S., Hollingsworth, T.N., Jean, M. & Mack, M.C. (2020) Factors shaping alternate successional trajectories in burned black spruce forests of Alaska. Ecosphere. 11 (5), e03129. doi:10.1002/ecs2.3129.

Jones, O.R., Barks, P., Stott, I., James, T.D., Levin, S., Petry, W.K., Capdevila, P., Che-Castaldo, J., Jackson, J., Römer, G., Schuette, C., Thomas, C.C. & Salguero-Gómez, R. (2022) Rcompadre and Rage—Two R packages to facilitate the use of the COMPADRE and COMADRE databases and calculation of life-history traits from matrix population models. Methods in Ecology and Evolution. 13 (4), 770–781. doi:10.1111/2041-210X.13792.

Jucker, T. (2022) Deciphering the fingerprint of disturbance on the three-dimensional structure of the world’s forests. New Phytologist. 233 (2), 612–617. doi:10.1111/nph.17729.

Jucker, T., Caspersen, J., Chave, J., Antin, C., Barbier, N., et al. (2017) Allometric equations for integrating remote sensing imagery into forest monitoring programmes. Global Change Biology. 23 (1), 177–190. doi:10.1111/gcb.13388.

Krause, C. & Lemay, A. (2023) Development and growth of young black spruce (Picea mariana) trees under two different hydrological conditions. Forest Ecology and Management. 541, 121083. doi:10.1016/j.foreco.2023.121083.

Kunstler, G., Guyennon, A., Ratcliffe, S., Rüger, N., Ruiz-Benito, P., Childs, D.Z., Dahlgren, J., Lehtonen, A., Thuiller, W., Wirth, C., Zavala, M.A. & Salguero-Gómez, R. (2021) Demographic performance of European tree species at their hot and cold climatic edges. Journal of Ecology. 109 (2), 1041–1054. doi:10.1111/1365-2745.13533.

Lloyd, A.H., Wilson, A.E., Fastie, C.L. & Landis, R.M. (2005) Population dynamics of black spruce and white spruce near the arctic tree line in the southern Brooks Range, Alaska. Canadian Journal of Forest Research. 35 (9), 2073–2081. doi:10.1139/x05-119.

Ma, Q., Su, Y., Niu, C., Ma, Q., Hu, T., Luo, X., Tai, X., Qiu, T., Zhang, Y., Bales, R.C., Liu, L., Kelly, M. & Guo, Q. (2023) Tree mortality during long-term droughts is lower in structurally complex forest stands. Nature Communications. 14 (1), 7467. doi:10.1038/s41467-023-43083-8.

MacDonald, G.M., Larsen, C.P.S., Szeicz, J.M. & Moser, K.A. (1991) The reconstruction of boreal forest fire history from lake sediments: A comparison of charcoal, pollen, sedimentological, and geochemical indices. Quaternary Science Reviews. 10 (1), 53–71. doi:10.1016/0277-3791(91)90030-X.

Maynard, D.S., Bialic-Murphy, L., Zohner, C.M., Averill, C., van den Hoogen, J., et al. (2022) Global relationships in tree functional traits. Nature Communications. 13 (1), 3185. doi:10.1038/s41467-022-30888-2.

McMahon, S.M., Parker, G.G. & Miller, D.R. (2010) Evidence for a recent increase in forest growth. Proceedings of the National Academy of Sciences. 107 (8), 3611–3615. doi:10.1073/pnas.0912376107.

Melvin, A.M., Mack, M.C., Johnstone, J.F., McGuire, A.D., Genet, H. & Schuur, E.A.G. (2015) Differences in ecosystem carbon distribution and nutrient cycling linked to forest tree species composition in a mid-successional boreal forest. Ecosystems. 18 (8), 1472–1488. doi:10.1007/s10021-015-9912-7.

Merow, C., Dahlgren, J.P., Metcalf, C.J.E., Childs, D.Z., Evans, M.E.K., Jongejans, E., Record, S., Rees, M., Salguero-Gómez, R. & McMahon, S.M. (2014) Advancing population ecology with integral projection models: a practical guide. Methods in Ecology and Evolution. 5 (2), 99–110. doi:10.1111/2041-210X.12146.

Metcalf, C.J.E., Horvitz, C.C., Tuljapurkar, S. & Clark, D.A. (2009) A time to grow and a time to die: a new way to analyze the dynamics of size, light, age, and death of tropical trees. Ecology. 90 (10), 2766–2778. doi:10.1890/08-1645.1.

Metcalf, C.J.E., McMahon, S.M., Salguero-Gómez, R. & Jongejans, E. (2013) IPMpack: an R package for integral projection models. Methods in Ecology and Evolution. 4 (2), 195–200. doi:10.1111/2041-210x.12001.

Mo, L., Zohner, C.M., Reich, P.B., Liang, J., de Miguel, S., et al. (2023) Integrated global assessment of the natural forest carbon potential. Nature. 624 (7990), 92–101. doi:10.1038/s41586-023-06723-z.

Morin, X., Fahse, L., Jactel, H., Scherer-Lorenzen, M., García-Valdés, R. & Bugmann, H. (2018) Long-term response of forest productivity to climate change is mostly driven by change in tree species composition. Scientific Reports. 8 (1), 5627. doi:10.1038/s41598-018-23763-y.

Morris, W.F. & Doak, D.F. (2002) *Quantitative conservation biology*lJ*: theory and practice of population viability analysis*. Sinauer Associates. https://cir.nii.ac.jp/crid/1130282270295869696.

Nappi, A., Drapeau, P., Saint-Germain, M. & Angers, V. (2010) Effect of fire severity on long-term occupancy of burned boreal conifer forests by saproxylic insects and wood-foraging birds. International Journal of Wildland Fire - INT J WILDLAND FIRE. 19. doi:10.1071/WF08109.

Nath, B. & Ni-Meister, W. (2021) The Interplay between Canopy Structure and Topography and Its Impacts on Seasonal Variations in Surface Reflectance Patterns in the Boreal Region of Alaska—Implications for Surface Radiation Budget. Remote Sensing. 13, 3108. doi:10.3390/rs13163108.

Needham, J., Merow, C., Butt, N., Malhi, Y., Marthews, T.R., Morecroft, M. & McMahon, S.M. (2016) Forest community response to invasive pathogens: the case of ash dieback in a British woodland. Journal of Ecology. 104 (2), 315–330. doi:10.1111/1365-2745.12545.

Needham, J., Merow, C., Chang-Yang, C.-H., Caswell, H. & McMahon, S.M. (2018) Inferring forest fate from demographic data: from vital rates to population dynamic models. Proceedings of the Royal Society B: Biological Sciences. 285 (1874), 20172050. doi:10.1098/rspb.2017.2050.

NEON (n.d.) Explore Field Sites | NSF NEON | Open Data to Understand our Ecosystems. https://www.neonscience.org/field-sites/explore-field-sites [Accessed: 31 May 2023].

Nicoll, B.C., Achim, A., Mochan, S. & Gardiner, B.A. (2005) Does steep terrain influence tree stability? A field investigation. Canadian Journal of Forest Research. 35 (10), 2360– 2367. doi:10.1139/x05-157.

Ohse, B., Compagnoni, A., Farrior, C.E., McMahon, S.M., Salguero-Gómez, R., Rüger, N. & Knight, T.M. (2023) Demographic synthesis for global tree species conservation. Trends in Ecology & Evolution. 38 (6), 579–590. doi:10.1016/j.tree.2023.01.013.

Palm, E.C., Suitor, M.J., Joly, K., Herriges, J.D., Kelly, A.P., Hervieux, D., Russell, K.L.M., Bentzen, T.W., Larter, N.C. & Hebblewhite, M. (2022) Increasing fire frequency and severity will increase habitat loss for a boreal forest indicator species. Ecological Applications. 32 (3), e2549. doi:10.1002/eap.2549.

Palviainen, M., Laurén, A., Pumpanen, J., Bergeron, Y., Bond-Lamberty, B., Larjavaara, M., Kashian, D.M., Köster, K., Prokushkin, A., Chen, H.Y.H., Seedre, M., Wardle, D.A., Gundale, M.J., Nilsson, M.-C., Wang, C. & Berninger, F. (2020) Decadal-Scale Recovery of Carbon Stocks After Wildfires Throughout the Boreal Forests. Global Biogeochemical Cycles. 34 (8), e2020GB006612. doi:10.1029/2020GB006612.

Pan, Y., Birdsey, R., Fang, J., Houghton, R., Kauppi, P., Kurz, W., Phillips, O., Shvidenko, A., Lewis, S., Canadell, J., Ciais, P., Jackson, R., Pacala, S., McGuire, A., Piao, S., Rautiainen, A., Sitch, S. & Hayes, D. (2011) A Large and Persistent Carbon Sink in the World’s Forests. *Science (New York*, N.Y.). 333, 988–993. doi:10.1126/science.1201609.

Pebesma, E. (2018) Simple Features for R: Standardized Support for Spatial Vector Data. The R Journal. 10 (1), 439–446.

Qi, M., Gadd, M., Martini, D.D., Davis, K., Xiong, B., Rosen, A., Hawes, N. & Salguero-Gomez, R. (2025) Biodiversity research requires more motors in the mud, air and water. https://ecoevorxiv.org/repository/view/8354/.

R Core Team (n.d.) R: A language and environment for statistical computing. R Foundation for Statistical Computing, Vienna, Australia. https://www.r-project.org/.

Reeb, D. (2024) Boreal forests: A global treasure. https://unece.org/sites/default/files/2025-03/2421978E_PDF_WEB.pdf.

Reid, K.A., Day, N.J., Alfaro-Sánchez, R., Johnstone, J.F., Cumming, S.G., Mack, M.C., Turetsky, M.R., Walker, X.J. & Baltzer, J.L. (2023) Black spruce (Picea mariana) seed availability and viability in boreal forests after large wildfires. Annals of Forest Science. 80 (1), 4. doi:10.1186/s13595-022-01166-4.

Roces-Díaz, J.V., Santín, C., Martínez-Vilalta, J. & Doerr, S.H. (2022) A global synthesis of fire effects on ecosystem services of forests and woodlands. Frontiers in Ecology and the Environment. 20 (3), 170–178. doi:10.1002/fee.2349.

Rosen, A., Battison, R., Hernández, C.M., Spacey, O., McLean, J., Prober, S., Gascoigne, S., McMahon, S., Jucker, T. & Salguero-Gómez, R. (2025) Modelling forest dynamics using integral projection models (IPMs) and repeat LiDAR.p.2025.01.06.631514. doi:10.1101/2025.01.06.631514.

Rotbarth, R., van Nes, E.H., Scheffer, M. & Holmgren, M. (2025) Boreal forests are heading for an open state. Proceedings of the National Academy of Sciences. 122 (2), e2404391121. doi:10.1073/pnas.2404391121.

Roussel, J.-R., Auty, D., Coops, N.C., Tompalski, P., Goodbody, T.R.H., Meador, A.S., Bourdon, J.-F., De Boissieu, F. & Achim, A. (2020) lidR: An R package for analysis of Airborne Laser Scanning (ALS) data. Remote Sensing of Environment. 251, 112061. doi:10.1016/j.rse.2020.112061.

Roussel, J.-R., documentation), D.A. (Reviews the, features), F.D.B. (Fixed bugs and improved catalog, segment_snags()), A.S.M. (Implemented wing2015() for, track_sensor()), B.J.-F. (Contributed to R. for, track_sensor()), G.D. (Implemented G. for, management), L.S. (Contributed to parallelization, code), S.A. (Author of the C. concaveman & function), B.S.-O. (Author of the ‘chm_prep’ (2024) *lidR: Airborne LiDAR Data Manipulation and Visualization for Forestry Applications*. https://cran.r-project.org/web/packages/lidR/index.html.

Rüger, N., Condit, R., Dent, D.H., DeWalt, S.J., Hubbell, S.P., Lichstein, J.W., Lopez, O.R., Wirth, C. & Farrior, C.E. (2020) Demographic trade-offs predict tropical forest dynamics. Science. doi:10.1126/science.aaz4797.

Salguero-Gómez, R. & Gamelon, M. (2021) Demographic Methods across the Tree of Life. Oxford, New York, Oxford University Press.

Salguero-Gómez, R., Jones, O.R., Archer, C.R., Buckley, Y.M., Che-Castaldo, J., et al. (2015) The compadre Plant Matrix Database: an open online repository for plant demography. Journal of Ecology. 103 (1), 202–218. doi:10.1111/1365-2745.12334.

Salguero-Gómez, R., Jones, O.R., Jongejans, E., Blomberg, S.P., Hodgson, D.J., Mbeau-Ache, C., Zuidema, P.A., de Kroon, H. & Buckley, Y.M. (2016) Fast–slow continuum and reproductive strategies structure plant life-history variation worldwide. Proceedings of the National Academy of Sciences. 113 (1), 230–235. doi:10.1073/pnas.1506215112.

Scholl, V.M., Cattau, M.E., Joseph, M.B. & Balch, J.K. (2020) Integrating National Ecological Observatory Network (NEON) Airborne Remote Sensing and In-Situ Data for Optimal Tree Species Classification. Remote Sensing. 12 (9), 1414. doi:10.3390/rs12091414.

Simpson, J.D. & Powell, G.R. (1981) Some Factors Influencing Cone Production on Young Black Spruce in New Brunswick. The Forestry Chronicle. 57 (6), 267–269. doi:10.5558/tfc57267-6.

Sturtevant, B.R. & Fortin, M.-J. (2021) Understanding and Modeling Forest Disturbance Interactions at the Landscape Level. Frontiers in Ecology and Evolution. 9. doi:10.3389/fevo.2021.653647.

Thorogood, R., Mustonen, V., Aleixo, A., Aphalo, P.J., Asiegbu, F.O., et al. (2023) Understanding and applying biological resilience, from genes to ecosystems. npj Biodiversity. 2 (1), 1–13. doi:10.1038/s44185-023-00022-6.

Tollefson, J. (2024) You’re not imagining it: extreme wildfires are now more common. Nature. doi:10.1038/d41586-024-02071-8.

Tsuyuzaki, S., Narita, K., Sawada, Y. & Kushida, K. (2014) The establishment patterns of tree seedlings are determined immediately after wildfire in a black spruce (Picea mariana) forest. Plant Ecology. 215 (3), 327–337. doi:10.1007/s11258-014-0303-5.

Van Cleve, K., Chapin, F.S., Flanagan, P.W., Viereck, L.A., Dyrness, C.T., Billings, W.D., Golley, F., Lange, O.L., Olson, J.S. & Remmert, H. (1986) *Forest Ecosystems in the Alaskan Taiga: A Synthesis of Structure and Function*. Ecological Studies. New York, NY, Springer. doi:10.1007/978-1-4612-4902-3.

Viereck, L.A., Werdin-Pfisterer, N.R., Adams, P.C. & Yoshikawa, K. (2008) Effect of wildfire and fireline construction on the annual depth of thaw in a black spruce permafrost forest in interior Alaska: a 36-year record of recovery. *In:* Kane, Douglas L.; Hinkel, Kenneth M.*, eds.* Proceedings of the Ninth International Conference on Permafrost*; June 29-July 3, 2008; Fairbanks, AK:* 1845–1850. https://research.fs.usda.gov/treesearch/32468.

Viglas, J.N., Brown, C.D. & Johnstone, J.F. (2013) Age and size effects on seed productivity of northern black spruce. Canadian Journal of Forest Research. 43 (6), 534–543. doi:10.1139/cjfr-2013-0022.

Viljur, M.-L., Abella, S.R., Adámek, M., Alencar, J.B.R., Barber, N.A., et al. (2022) The effect of natural disturbances on forest biodiversity: an ecological synthesis. Biological Reviews. 97 (5), 1930–1947. doi:10.1111/brv.12876.

Weinstein, B., Marconi, S., Zare, A., Bohlman, S., Graves, S., Singh, A. & White, E. (2020) *NEON Tree Crowns Dataset*. doi:10.5281/zenodo.3765872.

Weinstein, B.G., Marconi, S., Zare, A., Bohlman, S.A., Singh, A., Graves, S.J., Magee, L., Johnson, D.J., Record, S., Rubio, V.E., Swenson, N.G., Townsend, P., Veblen, T.T., Andrus, R.A. & White, E.P. (2024) Individual canopy tree species maps for the National Ecological Observatory Network.p.2023.10.25.563626. doi:10.1101/2023.10.25.563626.

Zambrano, J. & Salguero-Gómez, R. (2014) Forest Fragmentation Alters the Population Dynamics of a Late-successional Tropical Tree. Biotropica. 46 (5), 556–564. doi:10.1111/btp.12144.

Zuleta, D., Arellano, G., Muller-Landau, H.C., McMahon, S.M., Aguilar, S., Bunyavejchewin, S., Cárdenas, D., Chang-Yang, C.-H., Duque, A., Mitre, D., Nasardin, M., Pérez, R., Sun, I.-F., Yao, T.L. & Davies, S.J. (2022) Individual tree damage dominates mortality risk factors across six tropical forests. New Phytologist. 233 (2), 705–721. doi:10.1111/nph.17832.

