## Supplementary material for "LiDAR and hyperspectral-based structured population models show future forest fire frequency may compromise forest resilience": SOM

### 1.  Allometric model comparison

To determine which allometric model detects more of the larger tree crowns, we compared two window size allometric models against manual segmentation. We planned to only track the heights of individual trees with a crown area of at least 3 m^2^ because those trees heights would be informed by at least four CHM raster pixels. Individuals with a smaller crown area have high height variability with any misalignment of census scans (F. Fischer, pers. comm. 2023). We compared the number of 3 m^2^ individuals located manually to the number of individuals segmented by each window size allometric model. We completed the manual segmentation on a 150 m × 500 m strip that was entirely made up by *P. mariana* using QGIS (QGIS.org, 2024). The strip captured a dense to sparse canopy, as segmentation algorithms tend to struggle more on denser canopies (Battison et al., 2024; Aubry-Kientz et al., 2021). We set the minimum individual tree height to 4 m because the used LiDAR technology did not penetrate further down in the canopy (Dalponte & Coomes, 2016; Cao et al., 2023).

To choose the best window size allometric models, we compared both against the manual segmentation. To do so, we ran the segmentation algorithm on the CHM “highest” because the CHM is best for revealing treetops in the upper canopy, even with low point density LiDAR scans (Fischer et al., 2024). The first model was a log-log transformed linear allometric model, as often tree allometry models assume power-law behaviour in growth (Eq. S1, Huxley, Churchill & Strauss, 1993; Jucker et al., 2022). However, log-log linear allometric models tend to over-estimate tree crowns (Battison et al., 2024). Because of the possibility of over-segmentation with the log-log linear allometric model, we also considered a log-log transformed 95% quantile regression (Eq. S2). For both models (Eq. S1 and Eq. S2), we modelled crown diameter (D) as a function of height (H). In the log-log quantile regression (Eq. 2), $Q_{\tau=0.95}$ denotes the 95% percentile of the quantile to be a conditional distribution of crown diameter given height was modelled, which allows for a given height to have a wider range of possible crown areas (Battison et al. 2024). Both models were fit to a dataset of *P. mariana* individuals measured in the field at both of NEON’s interior Alaska sites, BONA and DEJU between 2016-2021 (NEON Vegetation Structure Data Product). If individuals had been measured multiple times, the first time both height and crown diameter was measured was included (Figure S1, Panel A for fitted models).

Log-log linear allometric model:

$$D \sim b*H^{a}$$

Eq. S1

Log-log quantile allometric model:

$$Q_{\tau=0.95}\left( H \right) \sim b*H^{a}$$

Eq. S2

The log-log quantile allometric model was a better input for the Dalponte and Coomes (2016) algorithm. The log-log quantile allometric model identified more trees over 3 m^2^ that overlapped with the manually segmented trees (211/863 > 73/863). In general, the log-log linear allometric model identified more treetops than the quantile regression, suggesting over-segmentation, as expected (2,945 > 1,238 treetops; Figure S1, Panel B for visualisation).

**Figure S1.** The allometric model comparison for height-dependent window size tree detection in the Dalponte and Coomes (2016) algorithm. Panel A: The log-log allometric models of height and crown diameter with the log-log linear regression (pink) and the log-log quantile regression (green) fitted to the field data of *P. mariana* at the BONA and DEJU site monitored by NEON. Panel B: The corresponding segmentation output with the log-log linear regression (pink, right to left hashing), the log-log quantile regression (green, left to right hashing) and the manual segmentation (blue).

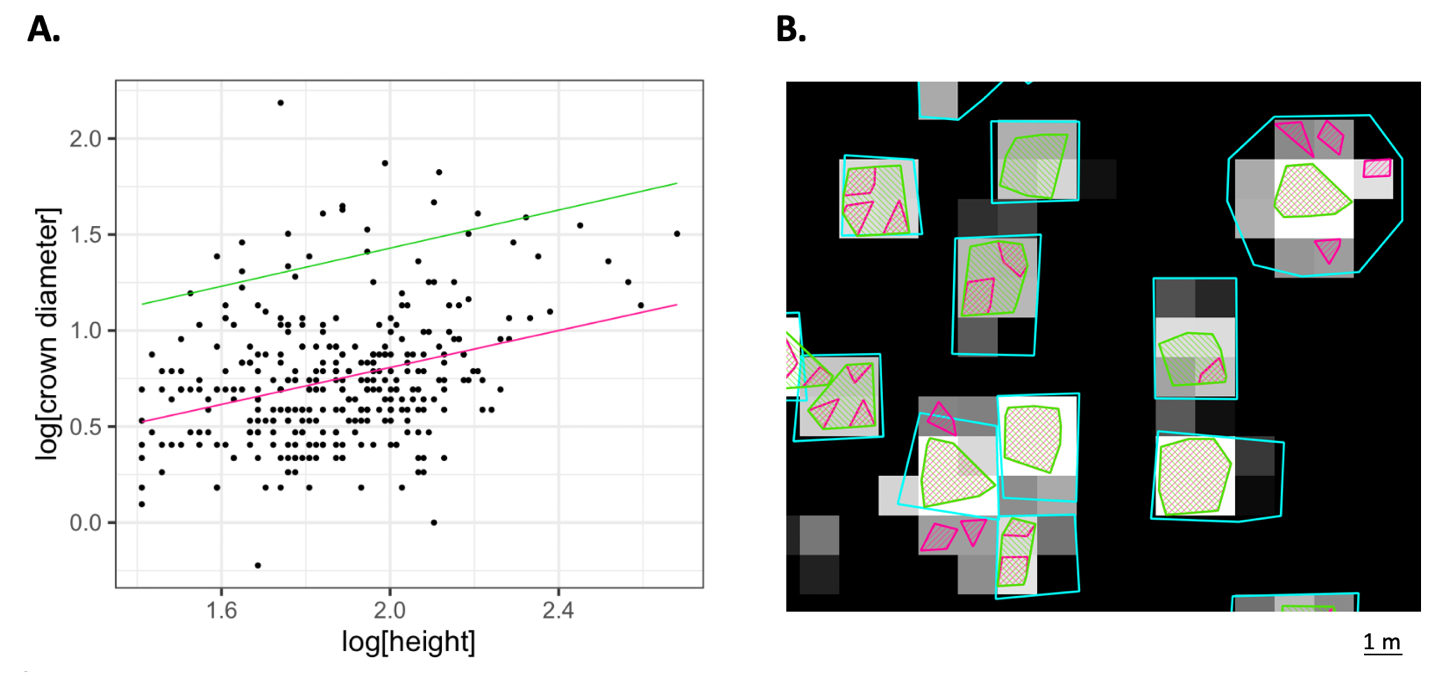

### 2. Tile selection criteria

We used a set of criteria for tile selection because only certain 1 km^2^ tiles fulfilled our selection criteria. First, we isolated the *P. mariana* stand within a 1999 fire patch, located on a North-facing slope (derived from satellite by Alaskan Wildland Fire Information Map Series, Figure 2B, tile 5). Our criterion for tile selection outside of the fire patch were tiles that were (1) dominated by *P. mariana*, (2) represented both North- and South-facing slopes, and (3) captured the elevation range of the site. Five tiles were sufficient to fulfil this criterion (Figure 2B, tiles 1-4 and 6). Single-species *P. mariana* stands were located using Weinstein et al.’s (2024) hyperspectral species map. The elevation map used is the digital terrain model (DTM) derived from LiDAR by (Fischer et al., 2024).

### 3. Trends in height change between censuses

**Figure S2:** We used the change in individual *P. mariana*’s height between 2017 and 2023 to characterise which trees survived (**A**) and how much they grew (**B**). **A**. The height change of the 42,816 individual *P. mariana* tree crowns tracked between 2017 and 2023 at BONA. Individuals in red decreased in height by 50% or more (*i.e.*, which here we assume as mortality; n = 345). **B**. The frequency of height changes in the trees that survived (Blue individuals in Panel A). The vertical dashed line indicates the mean growth (+ 0.35 m over the 6 year period).

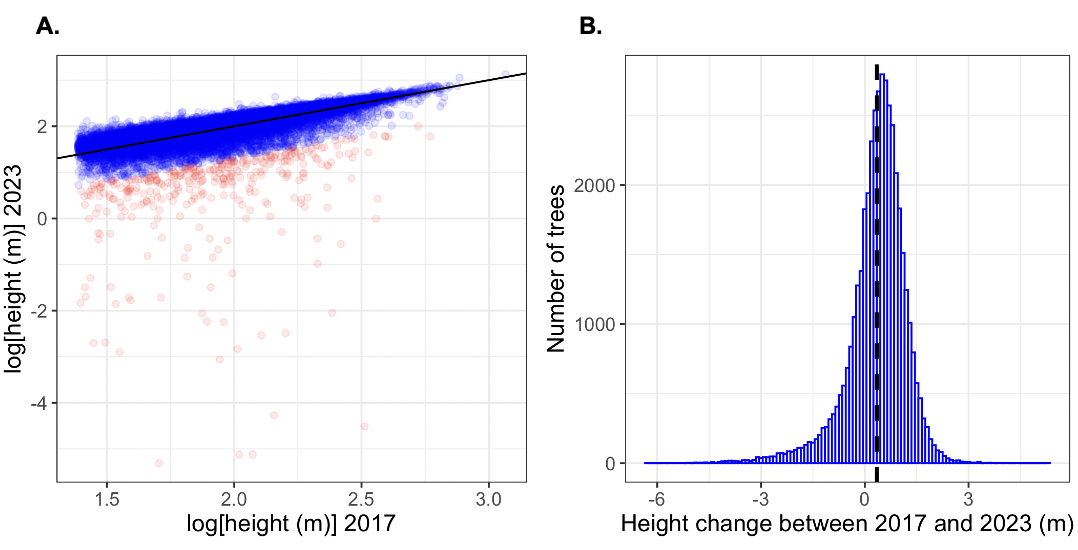

### 4. Surround canopy height sensitivity analysis

**Figure S3:** The estimates of passage time and mean life expectancy are insensitive to the choice of radius around the individual of interest. Panels A and C show estimates of passage time and life expectancy, respectively, when a 10 m radius is used. Similarly, Panels B and D show estimates of passage time and life expectancy, respectively, when a 20 m radius is used. The results are similar to the estimates with a 15 m radius, as used in the main study (Figure 3 and 4).

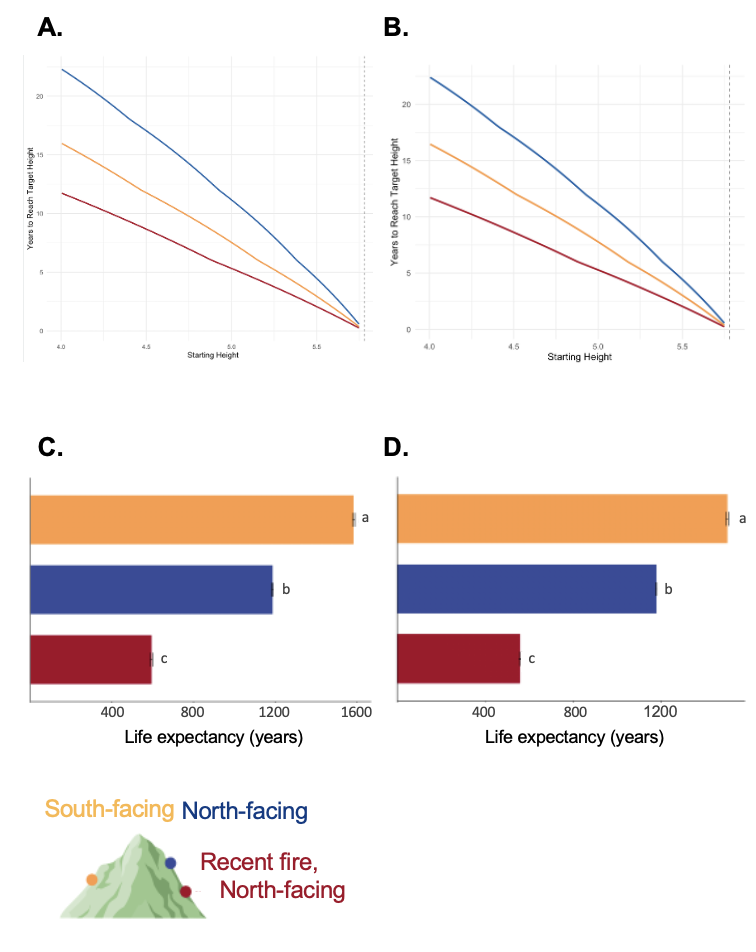

### 5. IPM Integration

We determined the lower ($L$) and upper ($U$) bounds of the $P$ kernel integral (Eq. 2) by subtracting or adding 10% of the minimum or maximum height observed in our dataset, respectively, a standard practice in IPMs (Easterling, Ellner & Dixon, 2000) . This approach amplifies the bounds of integration to help prevent ‘accidental eviction’ of individuals in the model (Williams, Miller & Ellner, 2012). This is an important step, as unintentional evictions can lead to the under-estimation of survival at large sizes (Easterling et al. 2000), and thus incorrect estimates of passage time and life expectancy. Thus, we determined which height transitions resulted in accidental evictions, and then overrode those transitions into a discrete class that represented the upper bound height *or higher*, following Williams, Miller & Ellner (2012). Lastly, we chose a 400×400 mesh size coupled with a mid-point integration of the *P* kernel to integrate our model, as commonly done in tree IPMs (Zuidema et al., 2010; Ellner, Childs & Rees, 2016; Needham et al., 2018). Choice of mesh size slightly higher or lower did not alter our estimates of passage time (Figure S4) and life expectancy (Figure S5) significantly.

**Figure S4:** The estimation of passage time is minimally sensitive to the choice of the number of meshpoints for IPM integration in any environmental conditions. The estimation of mean life expectancy is not sensitive to the choice of meshpoints for IPM integration in any environmental conditions. The chosen number of meshpoints for the main study was 400 meshpoints to balance accuracy with computational demands.

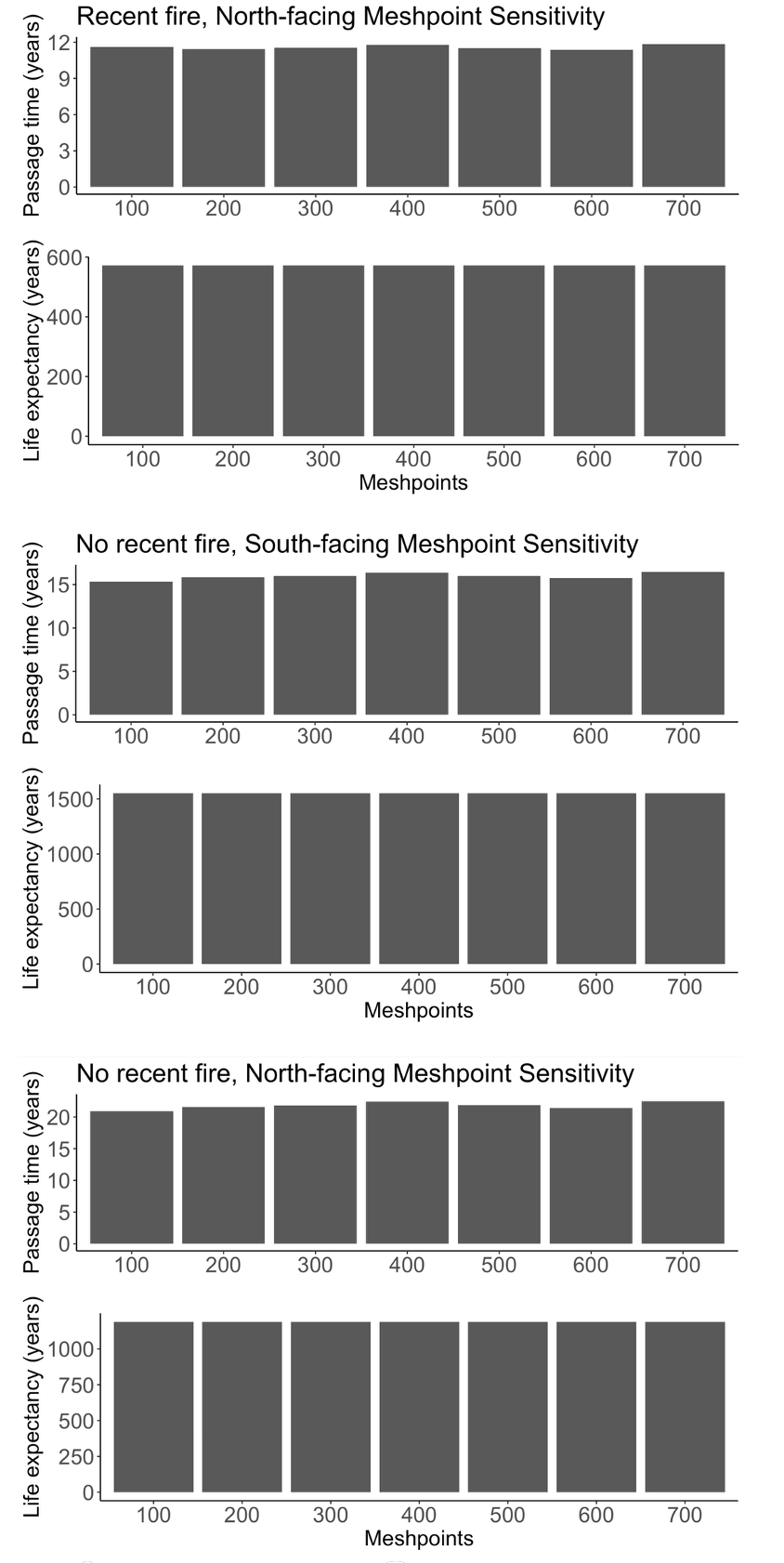

### 6. Description of environmental drivers

**Table S1**. A summary of the environmental drivers used to predict the survival and growth of *Picea mariana* populations between 2017 and 2023 in Alaska. The table contains information describing each variable, the source, and spatial resolution. DTM: Digital Terrain Model; CHM: Canopy Height Model.

| **Environmental driver** | **Description** | **Provenance** | **Resolution** |
| --- | --- | --- | --- |
| Elevation | Height above sea level (m) | 2017 DTM | 1 m^2^ |
| Slope steepness | Percentage change in DTM | 2017 DTM | 1 m^2^ |
| Aspect | On a north or south facing slope | 2017 DTM | 1 m^2^ |
| Surround canopy height (SCH) | Average height of the canopy within 15m radius of the tree | 2017 CHM | 1 m^2^ |
| Relative Change in Point density (RCP) | Point density 2023 – Point density 2017 / 2017 | Point density raster | 1 m^2^ |
| Recent fire | Whether the tree is growing in the 1999 fire patch or not. Non recent fire areas have not had a fire since 1940 (start of Map Series records) | Alaska Wildland Fire Information Map Series shapefile | 30 m^2^ |

### 7. Pearson correlation matrix

###### **Figure S5:** The Peason’s correlation matrix between all candidate predictor variables for survival and growth model. We calculated Pearson’s correlation coefficient (ρ) between all model predictors before constructing the survival and growth models with the aim to exclude one of the two variables involved in pair-wise correlations raising |ρ| > 0.6, following Beese et al. (2022). However, this step ended up not being necessary, as the highest correlation coefficient was below this threshold: between slope steepness and elevation (ρ = -0.56) and between surround canopy hight and height (ρ = 0.58).

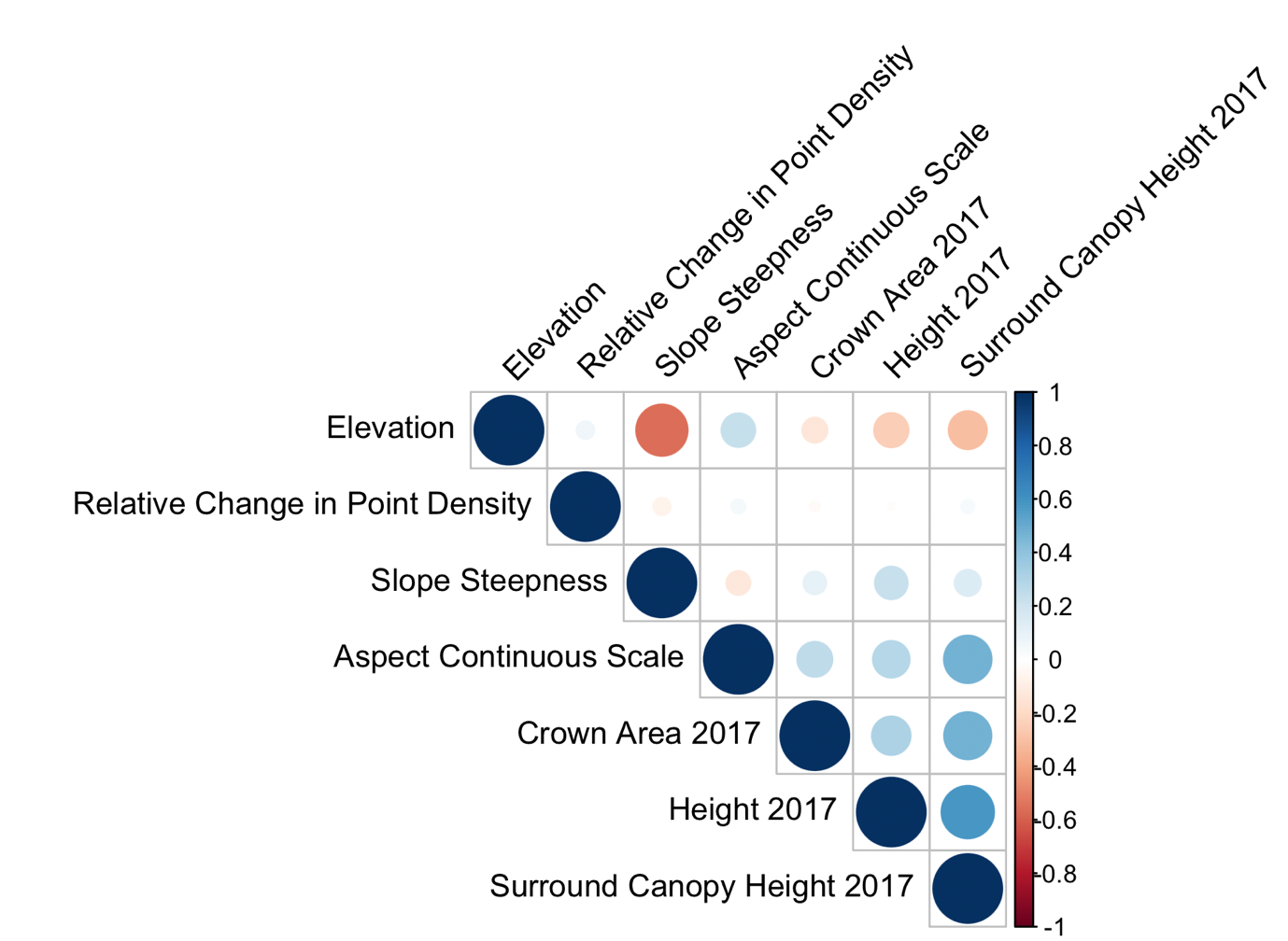

### 8. Inputs in the IPMs: mean values for environmental drivers

We quantified the impacts of key environmental drivers on survival and growth. The average elevation for an individual was 563.87 m, with similar number of individuals across the elevation range (180 – 820 m). A similar number of individuals were detected on North- and South-facing slopes (45.9% on North-facing slopes, 54.1% on South-facing slopes). A range of slope steepness was also covered from flat to steep (min = 0.59%, max = 39.96%). The average slope steepness was a gradual, reflecting most of the BONA *P. mariana* stands were on hill sides and ridges (mean = 12.27%). The average surround canopy height was less than the minimum tree height detected (3.16 m < 4 m), suggesting that the *P. mariana* stands are relatively sparse. As expected, the mean relative change in point density was positive (+3.8 RCP), reflecting the higher average point density across the forest in 2023 compared to 2017.

### 9. Survival: AIC comparison, model summary, and effect size

We used an automated stepwise Akaike Information Criterion (AIC) algorithm from the *R* package *MASS* to determine if any environmental driver brought about unnecessary complexity to our vital rate models (Venables & Ripley, 2002). For survival (Eq. 4), the most complex models had the lowest AIC score (>2 AIC score improvement), and as such we retained all environmental drivers in the model for the next steps of our analyses (See Figures S7-S5).

**Table S2.** The Akaike Information Criterion (AIC) algorithm chose the most complex model to predict survival from height and several a/biotic environmental variables. The table details how the AIC score is affected by removing each variable. Removing none of the variables resulted in the lowest AIC score by at least two points.

| **Removing variable** | **Df** | **Deviance** | **AIC** |
| --- | --- | --- | --- |
| none |  | 3365.9 | 3383.9 |
| Recent fire | 1 | 3372.8 | 3388.8 |
| Slope steepness | 1 | 3377.2 | 3393.2 |
| Elevation | 1 | 3380.9 | 3396.9 |
| Relative change in point density | 1 | 3382.5 | 3398.5 |
| North-facing | 1 | 3387.0 | 3403.0 |
| Log[Height 2017] | 1 | 3391.6 | 3407.6 |
| Log[Height 2017]^2^ | 1 | 3401.2 | 3417.2 |
| Surround canopy height | 1 | 3623.1 | 3639.1 |

**Table S3:** Summary of the final survival model (binomial logistic regression function of height, in log scale) that we integrated into the IPMs. For the no recent fire, South-facing slope IPM, the Norths term and Fires term aren’t added. For the no recent fire, North-facing slope IPM, the Fires aren’t added. Variables correspond to the variables in Eq. 4. The estimates are used in IPM integration to inform the directional effect each variable.

| **Variable** | **Description** | **Estimate** | **Std. Error** | **z value** | **P value** |
| --- | --- | --- | --- | --- | --- |
| $a_{s}$ | Intercept for North-facing slope and no recent fire | -6.717 | 2.243 | -2.994 | 0.003 |
| $b_{s}$ | Log height at t = 0 linear slope term | 11.927 | 2.215 | 5.384 | 7.29e-08 |
| $c_{s}$ | Log height at t = 0 quadratic slope term | -3.593 | 0.561 | -6.410 | 1.45e-10 |
| ${North}_{s}$ | Change in intercept for South-facing slope term | -0.841 | 0.191 | -4.416 | 1.00e-05 |
| ${Fire}_{s}$ | Change in intercept for recent fire term | 0.893 | 0.390 | 2.291 | 0.022 |
| ${Steepness}_{s}$ | Slope steepness slope term | -0.044 | 0.013 | -3.399 | 0.0007 |
| ${RCP}_{s}$ | Relative change in point  density slope term | 0.090 | 0.025 | 3.587 | 0.0003 |
| ${Elevation}_{s}$ | Elevation slope term | 0.002 | 0.0005 | 3.873 | 0.0001 |
| ${SCH}_{s}$ | Surround canopy height slope term | 0.820 | 0.059 | 13.977 | < 2e-16 |

**Table S4:** Effect size of environmental drivers and other model parameters on survival calculated as the standardised odds ratio and associated 95% confidence interval (CI). Calculated using the standardize_parameters function within the *effectsize* *R* package.

| **Parameter** | **Std Odds Ratio** | **95% CI** |
| --- | --- | --- |
| (Intercept) | 552.970 | [406.793, 772.823] |
| Log[height 2017] | 20.891 | [6.741, 61.765] |
| Log[Height 2017]^2^ | 0.028 | [0.010, 0.086] |
| North-facing | 0.431 | [0.294, 0.622] |
| Slope steepness | 0.816 | [0.726, 0.918] |
| Elevation | 1.296 | [1.137 1.477] |
| Recent fire | 2.443 | [1.227, 5.791] |
| Surround canopy height | 4.680 | [3.786, 5.838] |
| Relative change in Point Density | 1.422 | [1.186, 1.742] |

### 10. Growth: AIC comparison and model summary

**Table S5:** The Akaike Information Criterion (AIC) algorithm removed the quadratic term for height to predict growth from height and several a/biotic environmental variables. The final model is as follows: lm(formula = log_Ht_6 ~ log_Ht_0 + aspect_cat + slope_t0 + elevation_t0 + recentFire + canopy15_t0 + RCP).

| **Variable** | **Df** | **Sum of Sq** | **RSS** | **AIC** |
| --- | --- | --- | --- | --- |
| Log[Height 2017]^2^ | 1 | 0.004 | 601.58 | -180784 |
| None |  |  | 601.57 | -180782 |
| Slope steepness | 1 | 0.830 | 602.40 | -180726 |
| Relative change in point density | 1 | 1.874 | 603.44 | -180652 |
| Recent fire | 1 | 4.083 | 605.65 | -180497 |
| Elevation | 1 | 4.842 | 606.41 | -180444 |
| North-facing | 1 | 6.903 | 608.47 | -180300 |
| Log[Height 2017] | 1 | 10.950 | 612.52 | -180018 |
| Surround canopy height | 1 | 50.913 | 652.48 | -177334 |
| * Log[Height 2017]^2^ removed* | |  |  |  |
| **Variable** | **Df** | **Sum of Sq** | **RSS** | **AIC** |
| None |  |  | 601.58 | -180784 |
| Slope steepness | 1 | 0.83 | 602.40 | -180728 |
| Relative change in point density | 1 | 1.87 | 603.44 | -180654 |
| Recent fire | 1 | 4.08 | 605.66 | -180499 |
| Elevation | 1 | 4.95 | 606.53 | -180438 |
| North-facing | 1 | 7.02 | 608.60 | -180293 |
| Surround canopy height | 1 | 52.82 | 654.40 | -177212 |
| Log[Height 2017] | 1 | 1126.47 | 1728.04 | -135971 |

**Table S6:** Summary of the final growth model (ordinary least-squares regression function of height, in log scale) that we integrated into the IPMs. For the no recent fire, South-facing slope IPM, the Northg term and Fireg term aren’t added. For the no recent fire, North-facing slope IPM, the Fire aren’t added. Variables correspond to the variables detailed in Eq. 5. The estimates are used in IPM integration to inform the directional effect each variable.

| **Variable** | **Description** | **Estimate** | **Std error** | **t value** | **P value** |
| --- | --- | --- | --- | --- | --- |
| $a_{s}$ | Intercept for North-facing slope and no recent fire | 0.295 | 0.0073 | 40.514 | < 2e-16 |
| $b_{s}$ | Log height at t = 0 linear slope term | 0.800 | 0.003 | 281.981 | < 2e-16 |
| ${North}_{g}$ | Change in intercept for North-facing slope term | -0.0323 | 0.001 | -22.264 | < 2e-16 |
| ${Fire}_{g}$ | Change in intercept for recent fire term | 0.0786 | 0.005 | 16.980 | < 2e-16 |
| ${Steepness}_{g}$ | Slope steepness slope term | -0.001 | 0.0002 | -7.637 | 2.28e-14 |
| ${RCP}_{g}$ | Relative change in point density slope term | 0.002 | 0.0001 | 11.487 | < 2e-16 |
| ${Elevation}_{g}$ | Elevation slope term | 0.079 | 6.296e-06 | 18.692 | < 2e-16 |
| ${SCH}_{g}$ | Surround canopy height slope term | 0.027 | 0.0004 | 61.062 | < 2e-16 |

**Table S7:** Summary of the ANOVA for the model of growth, specifically to be able to compare the role of elevation, surround canopy height and slope steepness in growth for hypothesis H1.

| **Variable** | **Df** | **F value** | **P value** |  | **95% CI** |
| --- | --- | --- | --- | --- | --- |
| Log[Height 2017] | 1 | 160996.923 | < 0.001 | *** | [0.79, 1.00] |
| North-facing | 1 | 3733.624 | < 0.001 | *** | [0.08, 1.00] |
| Slope steepness | 1 | 141.412 | < 0.001 | *** | [2.47e-03, 1.00] |
| Elevation | 1 | 11.679 | < 0.001 | *** | [7.40e-05, 1.00] |
| Recent fire | 1 | 194.488 | < 0.001 | *** | [3.55e-03, 1.00] |
| Surround canopy height | 1 | 3823.953 | < 0.001 | *** | [0.08, 1.00] |
| Relative change in point density | 1 | 131.949 | < 0.001 | *** | [2.28e-03, 1.00] |
| Residuals | 42463 |  |  |  |  |

**Table S8:** The simplest model of growth, as summarised in this table, where future height is predicted only by past height, explains 76.12% (Adjusted R^2^) of the variance in future height. The model used in the IPMs, as summarised in Table S6, explains 79.92% (Adjusted R^2^) of the variance in future height. Therefore, adding environmental parameters to the model of growth explains an additional 3.8% of the variance in future height.

| **Term** | **Estimate** | **Std. Error** | **t value** | **Pr(>\|t\|)** |
| --- | --- | --- | --- | --- |
| Intercept | 0.216784 | 0.004753 | 45.61 | <2e-16 |
| Log Height 2017 | 0.910830 | 0.002475 | 367.96 | <2e-16 |

### 11. Growth variance: AIC comparison, model summary and ANOVA report

We quantified the error in the mean growth function $\varepsilon$ to capture the magnitude of uncertainty in LiDAR acquisitions (Clark, 2003). Specifically, we modelled the absolute value of the residuals in the growth function as a function of height at time $t$. To do so, we compared tree growth variation as constant and as linear function of height. We concluded, using AIC comparisons (Tables S9-10), that the linear function with height was the most parsimonious and explained the most variation in growth trajectories (*AICtab* function in the *AICcmodavg* *R* package; Mazerolle, 2023):

$$\left| \varepsilon\right|=a+bx \eta$$

Eq. S3

The variable $\eta$ in Eq. S3 represents unexplained variation in the absolute value of the growth function residuals.

We used an automated stepwise Akaike Information Criterion (AIC) algorithm from the *R* package *MASS* to determine if any environmental driver brought about unnecessary complexity to our vital rate models (Venables & Ripley, 2002). For the growth vital rate model (Eq. 5), we retained all environmental drivers, but removed the quadratic term for future height.

**Table S9:** A summary table of the AIC comparison between two possible models of growth variance around the mean. **“**growth_resid_mod1” modelled the absolute value of the residuals as a constant. “growth_resid_mod2” modelled variance as a function of log(height) in 2017. “growth_resid_mod2” has the lower AIC score by at least two points and was used in the IPMs.

| **Model** | **K** | **AICc** | **Delta_AICc** | **AICcWt** | **Cum.Wt** | **LL** |
| --- | --- | --- | --- | --- | --- | --- |
| Growth_resid_mod2 | 3 | -89675.32 | 0.00 | 1 | 1 | 44840.66 |
| Growth_resid_mod11 | 2 | -89238.64 | 436.68 | 0 | 1 | 44621.32 |

###### **Table S10:** A summary table of the coefficient estimates for the variables in the model of the growth variance around the mean. These estimates were used as the coefficients in the parameterisation of the IPMs as in Eq. 6.

| **Variable** | **Description** | **Estimate** | **Std error** | **t value** | **P value** |
| --- | --- | --- | --- | --- | --- |
| $a_{s}$ | Intercept for North facing slope and no recent fire | 0.148 | 0.003 | 47.96 | <0.001 |
| $b_{s}$ | Log height at t = 0 linear slope term | -0.03 | 0.002 | -21.00 | <0.001 |

### 12. Kernels with recent fire history and aspect combinations

**Figure S6:** IPM kernel prediction of the growth of a surviving individual of a given height in time t by time t +6 with different combinations of fire history and aspect conditions. The black points (high transparency) represent tree crown measurements between 2017 and 2023 at the BONA site that match the fire history and aspect of the respective IPM. **A**. The IPM with the combination of no recent fire and South-facing slope is plotted. **B**. The IPM with the combination of recent fire on a North-facing slope is plotted. **C**. The IPM with the combination of no recent fire on a North-facing slope is plotted. The final combination of recent fire history and South-facing is not included as there was no representative data. The greater intensity of the colour reflects the higher probability of a given transition from tree height $x$ to tree height $y$ between $t$ and $t+6$.

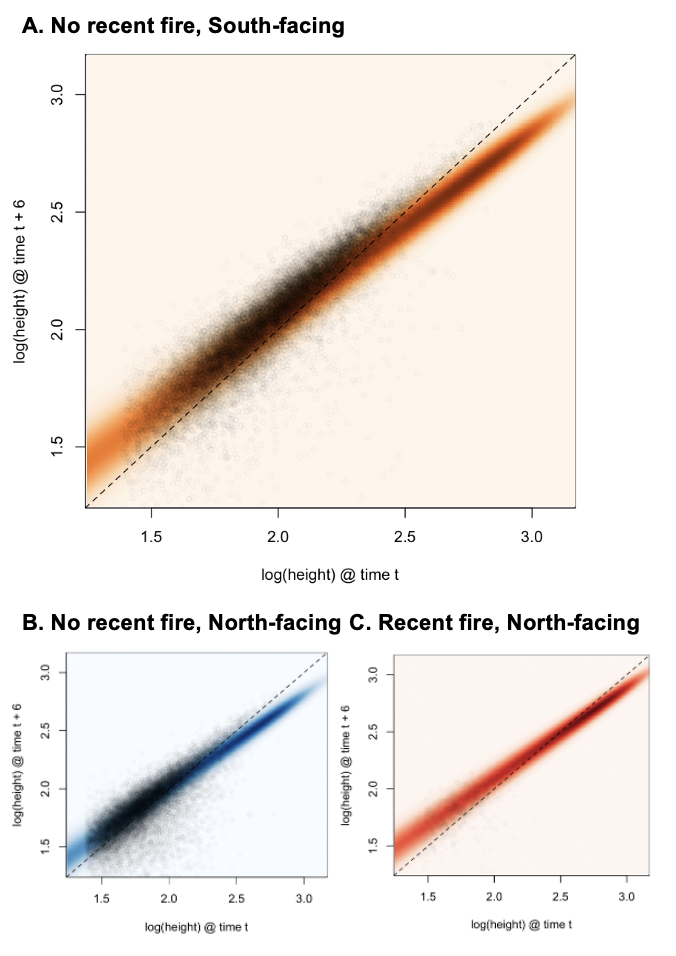

### 13. Distribution of tree heights in 1999 fire patch

**Figure S7:** The target height for the passage time analysis (5.78 m) is within the range of the heights within the 1999 fire patch. Here, the frequency of tree heights inside the 1999 fire patch in 2017 are shown, with a maximum height of 10.05 m. Tree heights are derived from extracting the maximum value of the Canopy Height Model (1 m^2^ resolution) within the segmented tree crown.

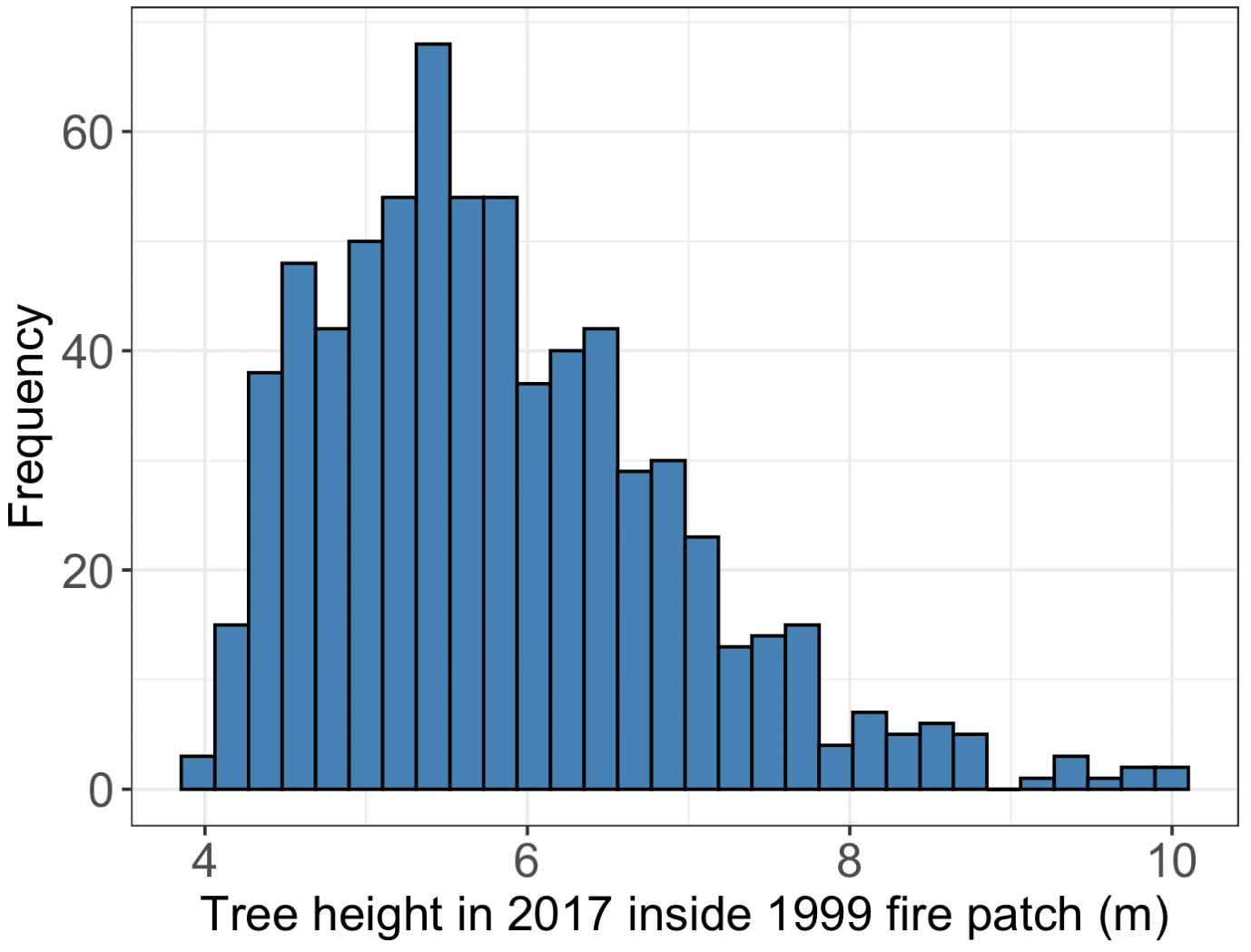

### 14. References

Aubry-Kientz, M., Laybros, A., Weinstein, B., Ball, J.G.C., Jackson, T., Coomes, D. & Vincent, G. (2021) Multisensor Data Fusion for Improved Segmentation of Individual Tree Crowns in Dense Tropical Forests. *IEEE Journal of Selected Topics in Applied Earth Observations and Remote Sensing*. 14, 3927–3936. doi:10.1109/JSTARS.2021.3069159.

Battison, R., Prober, S.M., Zdunic, K., Jackson, T.D., Fischer, F.J. & Jucker, T. (2024) *Tracking tree demography and forest dynamics at scale using remote sensing*.p.2024.06.11.598435. doi:10.1101/2024.06.11.598435.

Beese, L., Dalponte, M., Asner, G.P., Coomes, D.A. & Jucker, T. (2022) Using repeat airborne LiDAR to map the growth of individual oil palms in Malaysian Borneo during the 2015–16 El Niño. *International Journal of Applied Earth Observation and Geoinformation*. 115, 103117. doi:10.1016/j.jag.2022.103117.

Cao, Y., Ball, J.G.C., Coomes, D.A., Steinmeier, L., Knapp, N., Wilkes, P., Disney, M., Calders, K., Burt, A., Lin, Y. & Jackson, T.D. (2023) Benchmarking airborne laser scanning tree segmentation algorithms in broadleaf forests shows high accuracy only for canopy trees. *International Journal of Applied Earth Observation and Geoinformation*. 123, 103490. doi:10.1016/j.jag.2023.103490.

Dalponte, M. & Coomes, D.A. (2016) Tree-centric mapping of forest carbon density from airborne laser scanning and hyperspectral data. *Methods in Ecology and Evolution*. 7 (10), 1236–1245. doi:10.1111/2041-210X.12575.

Fischer, F.J., Jackson, T., Vincent, G. & Jucker, T. (2024) Robust characterisation of forest structure from airborne laser scanning—A systematic assessment and sample workflow for ecologists. *Methods in Ecology and Evolution*. n/a (n/a). doi:10.1111/2041-210X.14416.

Huxley, J.S., Churchill, F.B. & Strauss, R.E. (1993) *Problems of Relative Growth*. Johns Hopkins University Press. doi:10.56021/9780801846595.

Jucker, T., Fischer, F.J., Chave, J., Coomes, D.A., Caspersen, J., et al. (2022) Tallo: A global tree allometry and crown architecture database. *Global Change Biology*. 28 (17), 5254–5268. doi:10.1111/gcb.16302.
